# The CRAC channel inhibitor BTP2 restricts Tulane virus and human norovirus replication independent of store-operated calcium entry

**DOI:** 10.1101/2025.02.23.639787

**Authors:** Francesca J. Scribano, J. Thomas Gebert, Kristen A. Engevik, Nicole M. Hayes, Son Pham, Soni Kaundal, Janam J. Dave, B V Venkataram Prasad, Mary K. Estes, Sasirekha Ramani, Joseph M. Hyser

## Abstract

Human norovirus is the leading cause of viral gastroenteritis across all age groups. While there is a need for human norovirus antivirals, therapeutic development has been hindered by a lack of cell culture systems and animal models of infection. Surrogate viruses, such as Tulane virus (TV), have provided tractable systems to screen potential antiviral compounds. Our previous work demonstrated that Tulane virus encodes a viral ion channel, which dysregulates cytosolic calcium signaling. We set out to investigate whether host pathways triggered by viral ion channel activity, including store-operated calcium entry (SOCE), play a role in virus replication. Using pharmacologic inhibitors and genetically engineered cell lines, we establish that the SOCE inhibitor, BTP2, reduces TV replication in an SOCE-independent manner. We observed a significant reduction in TV replication, protein expression, and RNA synthesis in cells with both pre- and post-infection BTP2 treatment. By serial passage and plaque isolation, we demonstrate that TV quasi-species have mixed susceptibility and resistance to BTP2. Sequence comparison of the quasi-species revealed that amino acid changes in the structural proteins were associated with drug resistance. We utilized reverse genetics to generate TV with the resistance-associated VP1 and VP2 amino acid changes and found that a single amino acid change in VP1 (I380M) conferred BTP2 resistance. Further, expression of resistant VP2 alone was sufficient to partially rescue the replication of susceptible virus. Together, this supports that TV structural proteins are the targets of BTP2. Finally, using human intestinal organoids, we demonstrate that BTP2 significantly reduces human norovirus replication.

**Importance:** Our work identifies BTP2 as a potential human norovirus antiviral pharmacophore and highlights the utility of targeting calicivirus structural proteins to restrict viral replication. Further, we establish a system whereby Tulane virus can be used to screen novel antiviral candidates and establish their mechanism of action. Together, this will facilitate rapid preclinical validation of other novel human norovirus therapeutics.

## Introduction

Human norovirus (HuNoV) is the leading cause of non-bacterial gastroenteritis across all age groups [1, 2]. While HuNoV infection is acute and self-limiting in immunocompetent hosts, severe, chronic infections occur in vulnerable populations, including children, the elderly, and immunocompromised patients. In the United States, HuNoV infection accounts for 465,000 emergency department visits and 109,000 hospitalizations each year [3, 4]. Globally, HuNoV causes about 212,000 annual deaths [5]. To date, there are no approved prophylactic or therapeutic treatments that inhibit HuNoV replication or HuNoV-associated disease.

The development of HuNoV vaccines or antiviral therapies has been complicated by a limited understanding of the mechanisms of viral replication and pathogenesis. This is in large part due to historical challenges in propagating HuNoV in traditional cell culture systems and animal models. Currently, models for antiviral screening include surrogate virus systems such as Tulane virus (TV) or murine norovirus (MNV), the HuNoV replicon, and bona fide HuNoV replication in human intestinal organoids (HIOs) or zebrafish larvae. The recent finding that HuNoV can infect HIOs and zebrafish larvae has revolutionized the norovirus field [6–12]. However, while these systems productively replicate multiple GI and GII HuNoV genogroups [6, 7], they can be costly for high throughput antiviral screens or technically challenging to work with. Additionally, HIOs do not currently replicate HuNoVs with yields high enough to support continuous passage or production of virus stocks, so samples for virus inoculation are limited to patient stool. Therefore, utilizing surrogate viruses, including TV or MNV, can be useful to better understand mechanisms of replication, which can then be validated in more directed organoid studies with HuNoV, as has been done with virus disinfectant and inactivation studies [13–15].

TV, the prototype strain of the recovirus genus, was isolated from a rhesus macaque in 2008. Challenge studies suggest that TV-infected macaques present with a gastroenteritis-like disease, characterized by fever, diarrhea, duodenal inflammation, and virus shedding in the stool [16]. As new strains of recoviruses are identified, there is mounting evidence that some strains can be detected in human stool [17], neutralized by cross-reactive antibodies in human sera [18], and cultivated in continuous *in vitro* human cell lines [19]. Whether recoviruses contribute to the global burden of gastroenteritis in humans remains undetermined. However, recoviruses share many similarities with HuNoV and are a useful tool to investigate important aspects of norovirus biology. TV is a favorable surrogate model for HuNoV in that both viruses have similar genome organizations [20], bind to histo-blood group antigens [21], and cause acute diarrheal symptoms in permissive hosts [16]. Additionally, unlike HuNoV, TV can productively replicate in continuous monkey kidney cell lines, such as rhesus LLC-MK2 or vervet MA104 cells.

The utility of TV in the discovery of novel aspects of calicivirus infection is underscored by the recent finding that the nonstructural protein NS1-2 functions as an endoplasmic reticulum (ER)-localized viral ion channel (viroporin) [22, 23]. TV NS1-2 viroporin activity dysregulates cytosolic calcium signaling within infected cells, and transfection of HuNoV NS1-2 causes a similar pattern of calcium dysregulation [22]. Importantly, both ER and cytosolic calcium are important for TV replication, and we speculate that the same may be true for HuNoV [22]. While analogous viroporins, such as rotavirus NSP4, have been shown to promote virus replication both directly and via activation of host cell pathways [24–28], how dysregulation of intracellular calcium signaling aids in calicivirus replication remains unknown. In addition to directly affecting virus replication, viroporin activity engages host cell processes involved in maintaining calcium homeostasis, which independently may have important consequences for the virus life cycle. One such pathway is store-operated calcium entry (SOCE), which is triggered upon ER calcium store depletion. Two proteins, STIM and Orai, together are both necessary and sufficient for capacitive calcium entry through the SOCE pathway [29–33]. Previous studies have demonstrated that ER calcium release by rotavirus NSP4 activates SOCE and that inhibition of Orai or knockdown of STIM attenuates rotavirus replication [24, 26]. Given the importance of this pathway in enteric virus replication, and the likelihood of ER calcium release by calicivirus NS1-2 viroporin activity, we became interested in the role of SOCE in TV and HuNoV infection. Here we identify that the Orai channel inhibitor, BTP2, has antiviral activity against TV and two strains of HuNoV but demonstrate that this effect is SOCE-independent. Importantly, we also demonstrate that as the likely targets of BTP2, the calicivirus structural proteins may be attractive targets for further antiviral development. Together, this work underscores the utility of surrogate virus systems in the discovery and evaluation of novel candidates for the development of HuNoV therapeutics.

## Results

### Orai channel inhibitors block SOCE in MA104 cells but variably affect TV replication

We previously demonstrated that TV NS1-2 localizes to the ER and disrupts intracellular calcium signaling during infection in a manner similar to rotavirus NSP4, the prototype calcium-conducting viroporin [22]. Additionally, NSP4 triggers SOCE, and blocking this pathway *in vitro* significantly inhibits rotavirus replication [24]. Thus, we sought to determine the effect of several SOCE inhibitors on TV replication. We first confirmed that four well-characterized Orai channel blockers, BTP2, Ro2959, GSK-7975A (GSK), and Synta66 impair SOCE in MA104 cells by measuring cytosolic calcium concentrations during ER store depletion and calcium reintroduction [34]. In this assay, we measured relative cytosolic calcium by tracking the fluorescence of the genetically encoded calcium indicator, GCaMP6s. ER calcium stores were first depleted using thapsigargin to inhibit the sarcoplasmic/endoplasmic reticulum calcium ATPase (SERCA) pump in the presence of calcium-free medium (black arrow, Figure 1A). This enabled strong activation (i.e., opening) of SOCE channels in the plasma membrane without calcium influx into the cytoplasm. Calcium-containing media was then reintroduced, resulting in calcium entering the cell through the activated SOCE channels (red arrow, Figure 1A). By quantitating the maximum change in GCaMP6s fluorescence due to calcium influx, we found that all the compounds tested significantly reduced SOCE compared to the DMSO control (Figure 1B). Additionally, there were no cytotoxic effects of these inhibitors at the tested concentrations (Figure 1C). We next wanted to determine whether these inhibitors would impact TV replication. Although BTP2, Ro2959, GSK, and Synta66 all inhibited SOCE in MA104 cells, only BTP2 significantly reduced TV yield (Figure 1D). Given these discordant results, we next tested whether BTP2 inhibition of TV replication was independent of SOCE activity. Using CRISPR-Cas9, we developed a line of MA104 cells with a genetic knockout of STIM1, a protein required for SOCE activation [29, 30, 35]. This line showed loss of STIM1 protein expression by immunoblot (Supplemental Figure 1) and almost complete abrogation of SOCE-mediated calcium influx upon ER calcium depletion (Figure 1E-F). Together, these data validated the STIM1 knockout and showed the absence of compensatory mechanisms for SOCE in the absence of STIM1 in these cells. Despite the lack of SOCE, the STIM1 knockout did *not* affect TV replication, and BTP2 maintained its ability to significantly reduce TV yield (Figure 1G). This indicates that BTP2 blocks TV replication in an SOCE-independent manner. Given this, we sought to characterize BTP2-induced virus inhibition and to determine the stages of viral replication that were most significantly affected by compound treatment.

**Figure 1:**
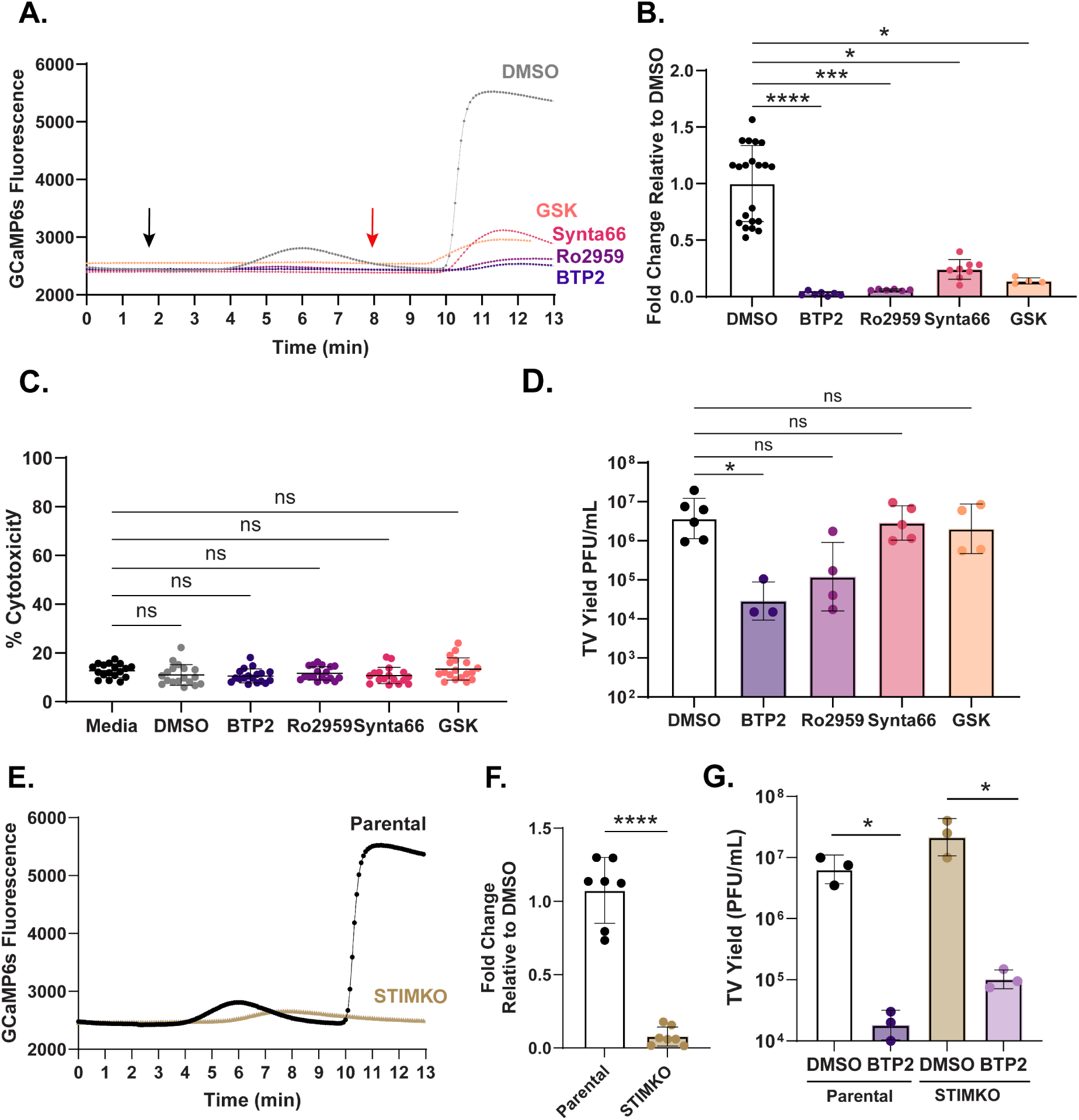
CRAC channel inhibitors block SOCE in MA104 cells but variably affect TV replication. **A**) Representative traces of GCaMP6s fluorescence following 1µM thapsigargin treatment (black arrow) and 2mM calcium perfusion (red arrow) in MA104G6s cells pretreated with DMSO control (grey), 5µM Ro2959 (purple) or 10µM BTP2 (blue), Synta66 (pink), or GSK (orange) for 20 min. **B**) Maximum fold change in GCaMP6s fluorescence per imaging field-of-view after 2mM calcium perfusion relative to DMSO control. All experiments were performed with a minimum of 3 biological repeats. **C**) LDH-based cytotoxicity in MA104G6s cells after 24-hr treatment with media alone (black), DMSO (grey) 10µM BTP2, Synta66, GSK or 5µM Ro2959. All experiments were performed with a minimum of 3 biological repeats of 6 technical replicates. **D**) TV yield in PFU/mL 24 hpi at MOI 1 with BTP2, Synta66, GSK (10µM) and Ro2959 (5µM). All compounds were added 1 hpi. Data are shown as an average of technical duplicates across at least 3 biological replicates. **E**) Representative traces of GCaMP6s fluorescence following thapsigargin treatment and 2mM calcium perfusion in parental MA104G6s cells (black) or STIM1 KO cells (gold). **F**) Maximum fold change in GCaMP6s fluorescence per imaging field-of-view after 2mM calcium perfusion relative to parental cells. Data are shown as a minimum of 2 technical replicates of at least 3 biological repeats. **G**) TV yield 24 hpi at MOI 1 in MA104G6s parental and STIM1 KO cells treated with DMSO control or 10µM BTP2 1 hpi. Data are shown as an average of technical duplicates across at least 3 biological replicates. For all experiments, normality was assessed by Shapiro-Wilk test. For SOCE and LDH experiments, ****p<0.0001 by Kruskal Wallis with Dunn’s multiple comparisons test after removal of outliers (ROUT Q=1%). TV yield assay data is plotted as geometric mean ± SD. *p<0.1 by Kruskal Wallis with Dunn’s multiple comparisons test or unpaired T test.

### BTP2 inhibits both early- and late-stage TV replication

To determine whether BTP2 inhibited other recovirus strains, we assessed the replication of FT285 and Mo/TG30 (GI.2 recoviruses) and FT7 (GI.3 recovirus) in the absence or presence of treatment. Interestingly, 10µM BTP2 did not affect FT7, FT285, or Mo/TG30 yield, demonstrating that BTP2 sensitivity was specific to TV (Supplemental Figure 2A-C). To better assess the effect of BTP2 on TV infection in MA104 cells, we first performed a dose range study and determined the 50% inhibitory concentration (IC_50_), which was achieved with a BTP2 concentration of ∼0.49µM (Figure 2A). Given that there was no additive inhibitory effect of BTP2 above 10µM (∼20X IC_50_), we used this concentration for our characterization studies, unless otherwise specified. To better characterize TV replication kinetics and BTP2 antiviral activity, we performed a time-course TV yield assay, adding BTP2 at 1-hour post-infection (hpi) and measuring virus yield at 0, 6, 12, and 24 hpi. BTP2 significantly reduced TV yield at 12 and 24 hpi (Figure 2B). To better define the stages of virus replication most significantly inhibited by compound treatment, we treated virus stock or cell monolayers with 10 µM BTP2 at specified time points before, during, or after infection (Figure 2C). Direct incubation of *TV stocks* with BTP2 for 1 hour did not affect the ability of the virus to infect and replicate compared to the DMSO control (Figure 2D), indicating that BTP2 did not neutralize virus particles. By contrast, incubating *cell monolayers* with BTP2 for 1 hour before or during TV inoculation did significantly reduce replication (Figure 2E-F). Likewise, adding BTP2 to the cell monolayers at 1 hpi and as late as 6 and 10 hpi significantly reduced virus yield (Figure 2G). Since these yield assays were performed at MOI 1, we further tested whether the effect of post-infection treatment might be due to inhibition of the early stages of secondary rounds of replication. Thus, we repeated the time of addition yield studies with an MOI 10 infection. These results mirrored our MOI 1 studies, in that BTP2 addition at 1, 6, and 10 hpi significantly reduced replication (Figure 2H), suggesting that the inhibitory effect occurs throughout the initial round of replication. This supports a model in which BTP2 treatment reduces replication kinetics in addition to inhibition of early events in the replication cycle. To determine if BTP2 impairs viral protein synthesis, we repeated the above yield assays and quantified TV VP1 and NS1-2 expression by western blot. Like the TV yield studies, BTP2 treatment prior to, during, or at 1, 6, or 10 hours post inoculation reduced TV protein expression (Figure 3A, B). Our western blot data also showed that viral protein synthesis remained high by 10 hpi, providing further insight into why BTP2 treatment at this later time point still restricts viral replication (Figure 3A, B). To determine the effect of BTP2 on genome replication, we measured TV genome equivalents at 2 and 24 hpi with the same BTP2 treatment schemes. Paralleling our yield and protein expression data, pre- and post-infection BTP2 treatment significantly reduced levels of TV RNA at 24 hpi and genome replication between 2 and 24 hpi (Figure 3C, D). Together, this data suggests that BTP2 inhibits the viral life cycle at a point prior to translation and genome replication.

**Figure 2:**
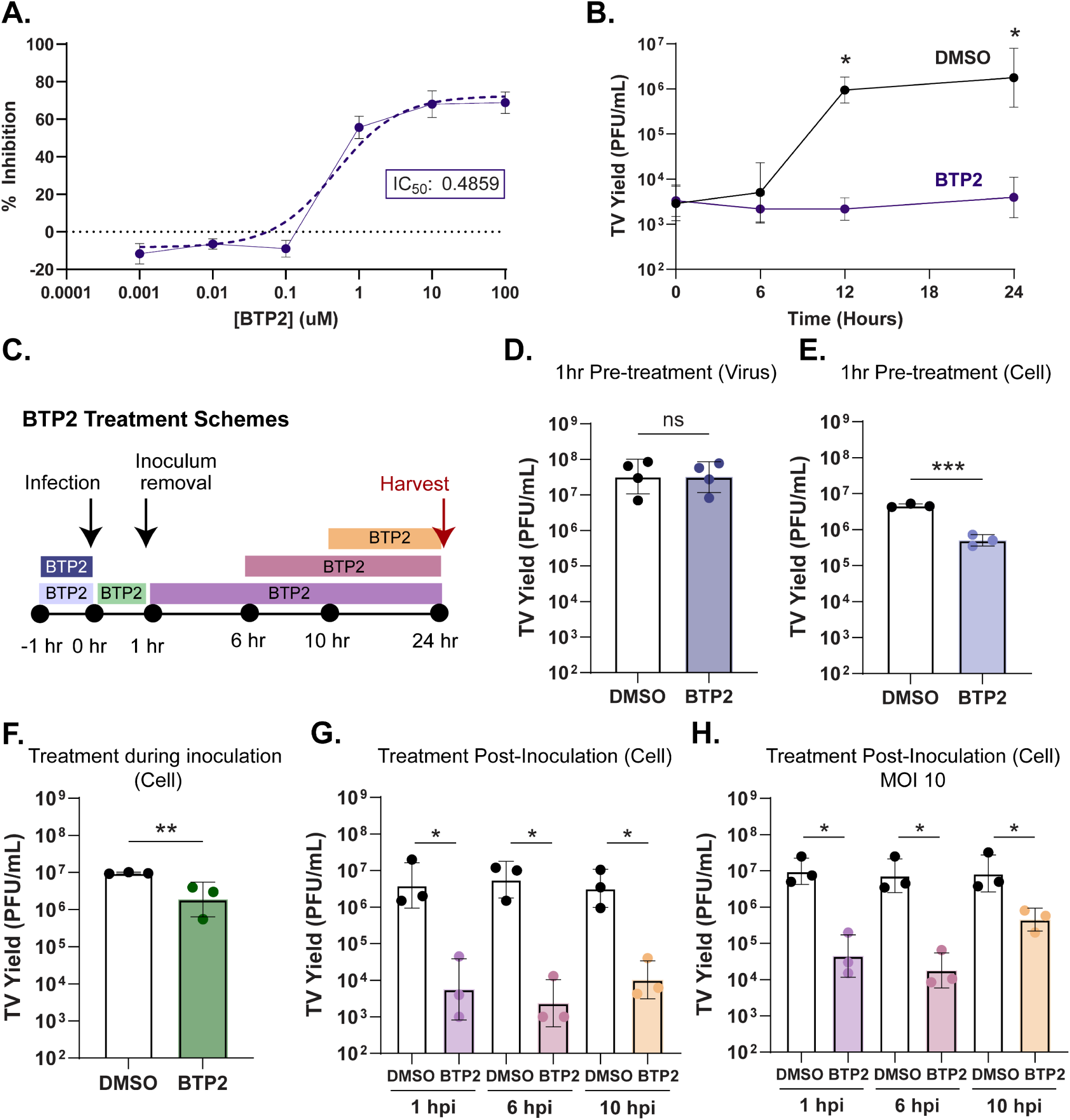
BTP2 inhibits both early- and late-stage TV replication. **A**) Percent inhibition of monolayer clearance in crystal violet-based assay in TV infected monolayers (MOI 10) treated with 0.001, 0.01, 0.1, 1, 10, or 100µM BTP2. Dose curve was fit with a non-linear regression (dotted line) to estimate the inhibitory concentration 50 (IC50). **B**) Time course assay measuring TV yield in PFU/mL at 0, 6, 12, and 24 hpi with DMSO (black) or BTP2 (purple) treatment at 1hpi with an MOI 1 infection. **C**) Schematic of BTP2 time-of-treatment. 10µM BTP2 stock was incubated with TV stock or cell monolayers for 1 hour prior to infection, or with cell monolayers during the 1 hr inoculum incubation or 1, 6, or 10 hours post inoculation. All infections were harvested at 24 hpi. **D-G**) plaque assay titrations of TV yield following MOI 1 infection and treatment with the corresponding pre-treatment scheme of TV stock (D) or pre- and post-treatment schemes of cell monolayers (E-G). **H**) Plaque assay titration of TV yield after MOI 10 infection and treatment with BTP2 at 1, 6, or 10 hpi. All data are shown as an average of technical duplicates across at least 3 biological replicates. For all experiments, normality was assessed by Shapiro-Wilk test. TV yield assay data is plotted as geometric mean ± SD. **p<0.01 by Kruskal Wallis with Dunn’s multiple comparisons test or unpaired T test.

**Figure 3:**
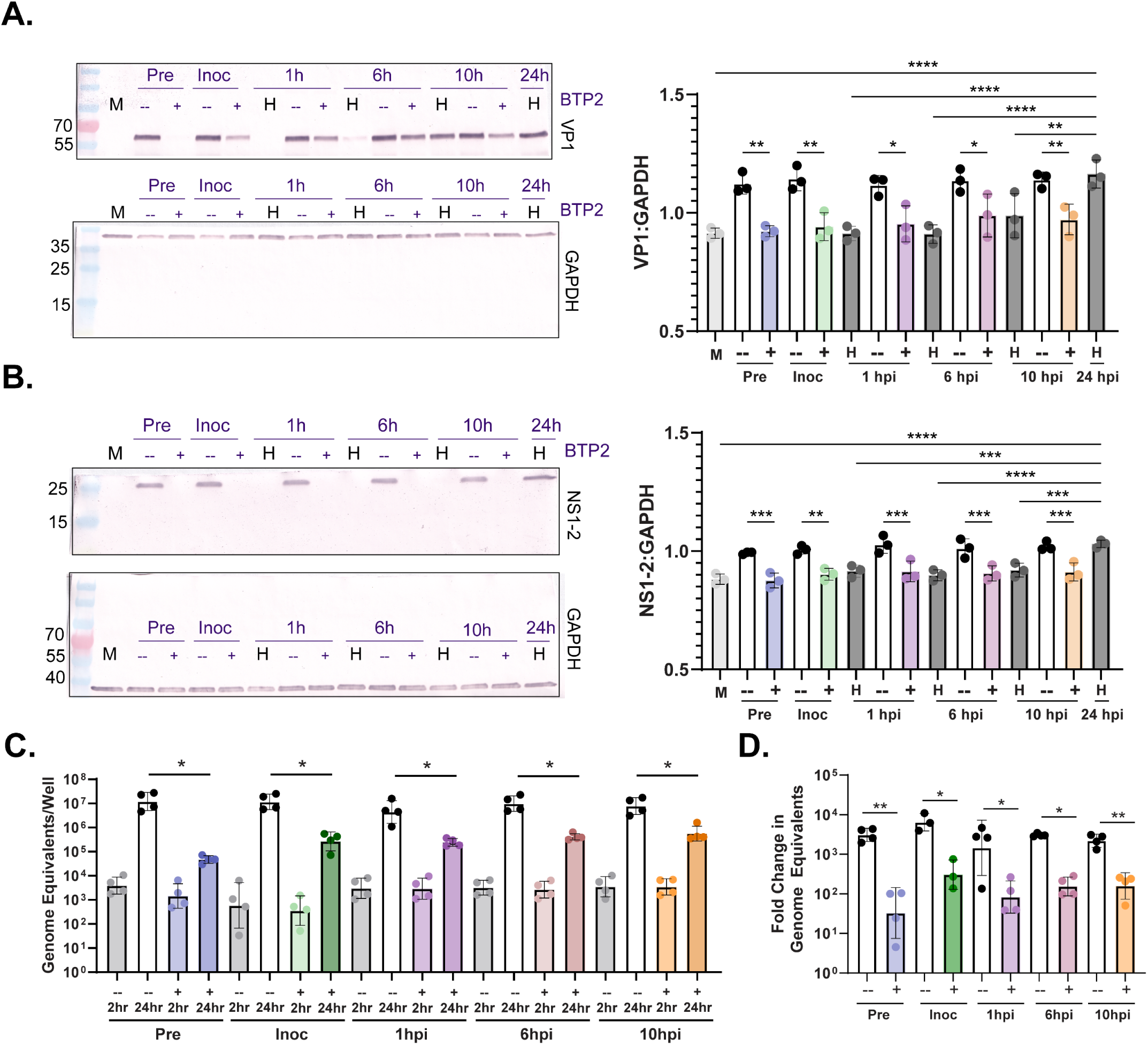
Early and late stage BTP2 addition reduces TV protein and RNA synthesis. **A-B**) TV VP1 protein expression (A) or NS1-2 expression (B) detected by Western blot following DMSO (-) or 10µM BTP2 (+) addition at 1 hour before infection (pre), during viral inoculation (inoc), or 1, 6, or 10 hrs post infection at an MOI of 1. In parallel infections, cell lysates from an untreated infection were harvested (H) at the indicated time points. GAPDH (bottom) served as the loading control. For quantitation, band intensity was plotted relative to GAPDH. Each point represents an independent biological replicate. **C**) TV genome equivalents determined through RT-qPCR of infected cells treated with DMSO (-) or BTP2 (+) at the indicated time points. RNA was harvested at 2 and 24 hpi, and genome equivalents were determined based on a standard curve from in vitro transcribed TV RNA. **D**) Fold change in TV genome equivalents between 2 and 24 hrs with DMSO (-) or BTP2 (+) treatment at the indicated time points. Data are shown from four biological replicates, and the RT-qPCR data represents an average of technical duplicates for each biological replicate. ****p<0.0001 by 1-way ANOVA with Sidak’s multiple comparisons test or unpaired T test.

### TV overcomes BTP2 susceptibility with passage through emergence of a resistant variant

The time-of-addition experiments suggested that BTP2 treatment inhibits pre-transcriptional and/or translational phases of the viral life cycle; however, they did not allow for the determination of whether specific viral proteins are inhibited. To determine if there was a viral target of BTP2, we serially passaged TV in the presence of BTP2 to rescue a BTP2-resistant mutant. We used crystal violet to stain infected cell monolayers and observed that DMSO-passaged virus, at every passage, caused complete cytopathic effect (CPE) when treated with the DMSO vehicle control. In contrast, in the presence of BTP2, DMSO-passaged virus caused little CPE, suggesting that BTP2 protected cells from TV-induced cell death (Figure 4A). Conversely, BTP2-passaged TV caused CPE when treated with DMSO but became increasingly resistant to BTP2-mediated monolayer protection with each passage (Figure 4B). We confirmed by virus yield assay, using passage 4 virus, that the DMSO-passaged virus maintained its BTP2 susceptibility, while the BTP2-passaged virus was BTP2 resistant (Figure 4C). We considered two possibilities for the isolation of BTP2-resistant TV: it may have arisen either by *de novo* mutation or by selection of naturally resistant quasi-species in our virus stock. Since BTP2 resistance emerged after only four passages in culture, we suspected the latter. To test this, we performed plaque assays with *non-drug passaged* virus stock, adding BTP2 or DMSO control to the overlay. In the presence of BTP2, TV plaques were still observed at dilutions of 10^-3^ or lower (Figure 4D), but in the presence of the DMSO vehicle, plaques were observed out to the 10^-6^ dilution. Additionally, the plaques that were formed from both treatments showed no difference in plaque size (Figure 4D). Together, these data indicate that the virus that initiated the plaques in each respective treatment had no defect in multi-round viral spread. By extension, BTP2 resistant virus was present in our non-passaged stocks, although at far lower concentrations than the BTP2 susceptible virus. To confirm the presence of BTP2 resistant virus in our non-passaged stocks, we isolated virus from the recovered plaques and found that the plaque picks from wells with BTP2-containing overlay were BTP2 resistant, while plaque picks from wells with DMSO-containing overlay were BTP2 susceptible (Figure 4E-F). These results confirmed that our starting viral stock contained a mixture of BTP2 susceptible and resistant TV quasi-species.

**Figure 4:**
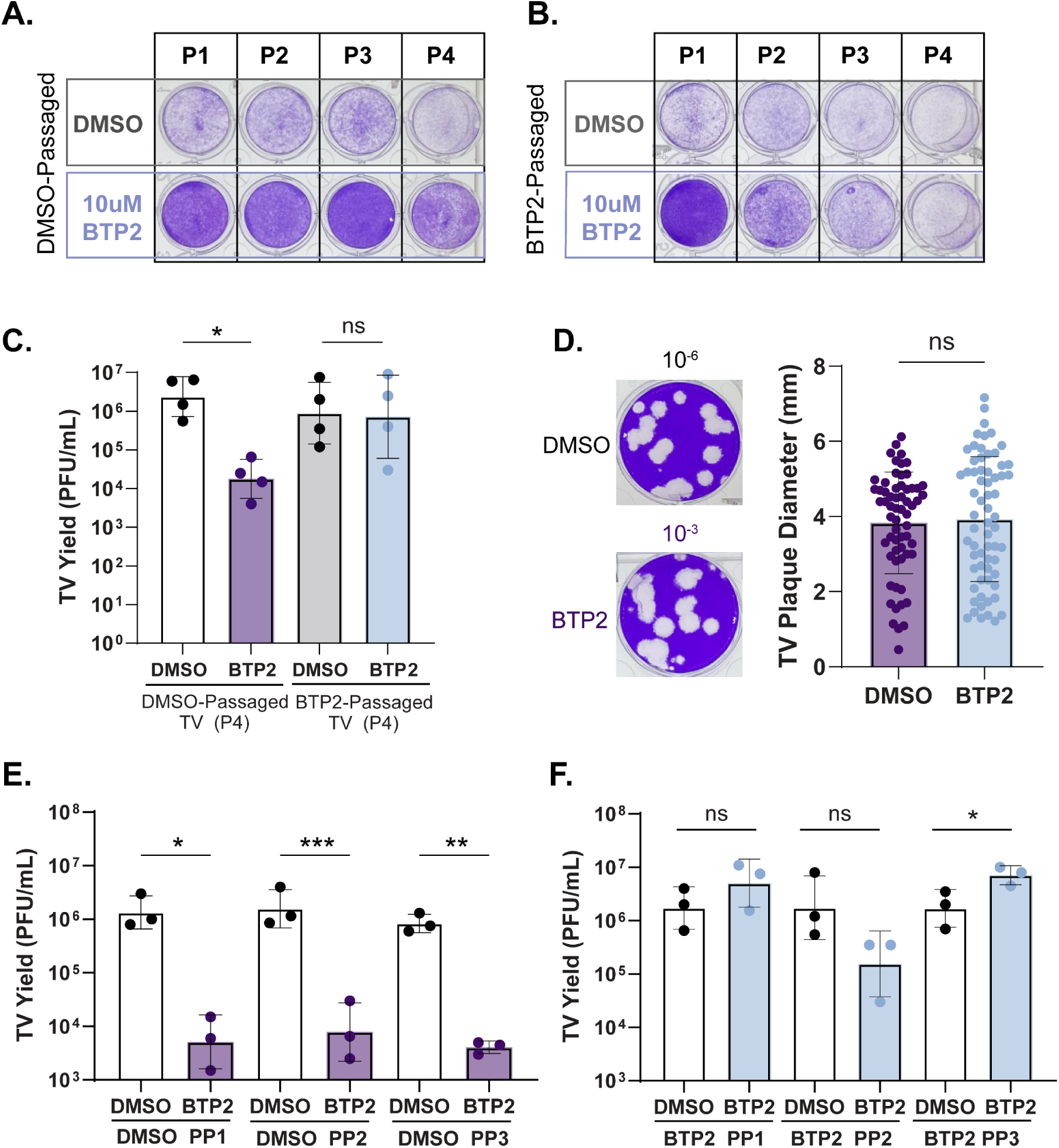
TV overcomes BTP2 susceptibility with passage through emergence of resistant quasi-species. **A**) Representative images of crystal-violet stained MA104 cell monolayers after infection with DMSO-passaged TV (A, passage (p) 1-4) or BTP2-passaged TV (B, passage (p) 1-4) in the presence of DMSO or 10µM BTP2. **C**) TV yield in PFU/mL from passage 4 DMSO or BTP2 passaged virus, challenged with DMSO (white, grey) or 10µM BTP2 (purple, light blue). Data are shown as an average of technical duplicates across at least 3 biological replicates. **D**) Representative images of crystal violet-stained plaques following DMSO or 10µM BTP2 addition to the plaque assay overlay (left panel) and quantitation of plaque diameter (millimeters) across three independent infections (right panel, n>62 plaques). **E-F**) TV yield in PFU/mL from DMSO (E) or BTP2 (F) solid overlay plaque picks, challenged with DMSO (white) or 10µM BTP2 (purple). Data are shown from at least 3 biological replicates. For all experiments, normality was assessed by Shapiro-Wilk test. TV yield assay data is plotted as geometric mean ± SD. **p<0.01 by Mann-Whitney or unpaired T test. Plaque diameter data was analyzed by Mann-Whitney U test.

### TV structural proteins mediate BTP2 susceptibility

The BTP2 plaque isolation experiments allowed us to distinguish multiple BTP2 susceptible and resistant TV variants, and we sought to determine whether they contained resistance-associated amino acid differences. To isolate clonal populations of susceptible and resistant virus, we performed three rounds of serial plaque isolation in the absence or presence of BTP2. We utilized virus inhibition assays to determine the degree of susceptibility of each of the clones to BTP2 treatment (Figure 5A). We noted that one of the BTP2 isolated clones, resistant clone 3, had an intermediate susceptibility phenotype (Figure 5A). We next performed yield assays with one representative susceptible and one representative resistant clone to verify that the BTP2 serial plaque isolation yielded BTP2 resistant virus, while the DMSO serial plaque isolation yielded BTP2 susceptible virus (Figure 5B). To determine if BTP2 resistance conferred a fitness cost or benefit, we performed growth curves using the same representative resistant and susceptible isolate. We found that BTP2 susceptibility does not affect the ability of these viruses to grow in cell culture in the absence of treatment (Figure 5C). Given the distinct effects of BTP2 treatment on these isolated viral variants, we utilized Sanger sequencing to identify sequence differences associated with resistance. In total, we detected one amino acid difference in VP1 (I380M), which was universally represented in all of the BTP2 resistant clones and a cluster of non-uniform amino acid differences at the C-terminus of VP2, which were present in two of the resistant clones (clone 1: K148N, I191T clone 2: Y156H, S179T) (Figure 5D, Supplemental Figure 3-4). No conserved amino acid changes were found in the ORF1 nonstructural proteins in any of the clones. This finding is consistent with cell-free functional assays that showed that BTP2 treatment did not affect TV or HuNoV protease activity (Supplemental Figure 5A-B), or HuNoV polymerase activity (Supplemental Figure 5C). To validate the role of the structural proteins in BTP2 resistance, we utilized reverse genetics to rescue TV with the resistance-associated amino acid changes in VP1 only, VP2 only, or both VP1 and VP2. The susceptible and resistance-associated sequences were generated from susceptible clone 1 and resistant clone 1 (Figure 5E, Supplemental Figure 6A, 7A). As expected, the rescued TV with BTP2 susceptible-associated VP1 and VP2 showed significantly reduced yield with BTP2 treatment (Figure 5F). Recombinant TV with the VP2 only resistance-associated amino acid changes also demonstrated susceptibility to BTP2 treatment. However, the viruses with the VP1 only and dual VP1, VP2 resistance-associated amino acid changes were BTP2 resistant (Figure 5F). There were no significant differences in BTP2 susceptibility across a range of doses between the VP1 only and combination VP1, VP2 viruses or the susceptible and VP2 only viruses (Figure 5G). Together, this suggests that the VP1 I380M mutation is the main driver of BTP2 resistance and a likely target of compound treatment. Mapping of the I(380) residue onto the TV VP1 cryo-EM structure shows that it is positioned at the VP1 dimer interface (Supplemental Figure 6B).

**Figure 5:**
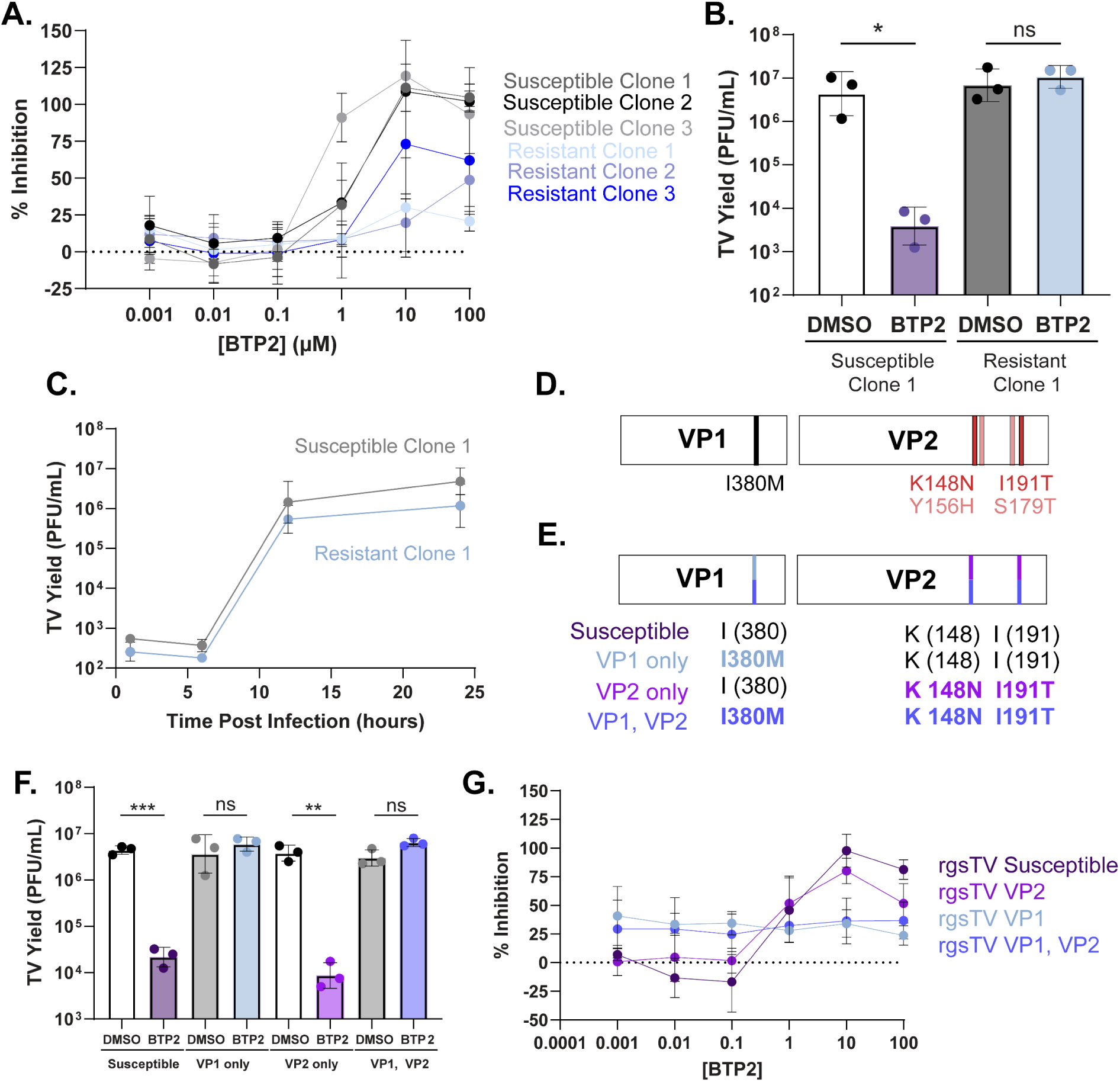
TV structural proteins mediate BTP2 susceptibility. **A**) Percent inhibition of monolayer clearance in crystal violet-based assay with 3 serial plaque isolated susceptible viruses (grey) and 3 serial isolated resistant viruses (blue) (MOI 3) treated with a dose range of BTP2. **B**) TV yield assay with one serial plaque isolated susceptible virus (clone 1) and one serial isolated resistant virus (clone 1) challenged with DMSO or 10µM BTP2. **C**)Time course assay measuring yield at 0, 6, 12, and 24 hpi with one serial plaque isolated susceptible virus (clone 1, grey) and one serial isolated resistant virus (clone 1, blue). **D**) Schematic of amino acid differences identified between BTP2 susceptible and resistant clones. **E**) Schematic of susceptible or resistance-associated amino acids in the four reverse genetics Tulane viruses. **F**) Reverse genetics TV yield at 24hpi following an MOI 1 infection with DMSO or 10µM BTP2 treatment at 1hpi. **G**) Percent inhibition of monolayer clearance with reverse genetics TV (MOI 3) treated with BTP2 at the indicated dose range. For yield experiments, normality was assessed by Shapiro-Wilk test and data is plotted as geometric mean ± SD. ***p<0.001 by unpaired T test.

### Resistant VP2 partially rescues the replication of BTP2 susceptible TV

Given that the amino acid differences in VP2 were non-uniform and were not sufficient to rescue TV replication in the presence of BTP2 we next sought to validate whether VP2 played any role in mediating BTP2 resistance. To do this, we performed co-transduction and infection studies and determined if overexpression of the resistant VP2 was sufficient to confer BTP2 resistance to the plaque-isolated BTP2-susceptible virus clone. We generated adeno-associated viral (AAV) vectors for recombinant expression of VP2 from either susceptible or resistant clone 1. The VP2 from these constructs has an N-terminal 3xFLAG-tag and expressed mCherry downstream of an encephalomyocarditis virus internal ribosome entry site to aid in quantitating protein expression and transduction efficiency. We utilized an AAV vector expressing only the fluorescent protein, mScarlet, as an irrelevant protein overexpression control. We first validated AAV transduction efficiency by measuring mScarlet or mCherry fluorescence and FLAG by western blot (Figure 6A-D), which confirmed that the susceptible and resistant VP2 constructs were expressed to similar levels. Next, we determined the yield of a BTP2 susceptible clone of TV following infection of mScarlet or VP2-transduced monolayers treated with DMSO or BTP2. We found that in all cases BTP2 treatment significantly reduced viral yield (Figure 6E). We next analyzed the fold reduction in TV replication between DMSO and BTP2 treatment for each transduction condition. We found that the degree of BTP2 inhibition was not statistically different between monolayers expressing mScarlet or the susceptible VP2 or between susceptible VP2 and resistant VP2 expression. However, BTP2 inhibition was significantly attenuated in monolayers expressing the resistant VP2 compared to mScarlet alone (Figure 6F), suggesting that overexpression of VP2 from the BTP2-resistant clone may confer partial, but not complete resistance to BTP2 treatment.

**Figure 6:**
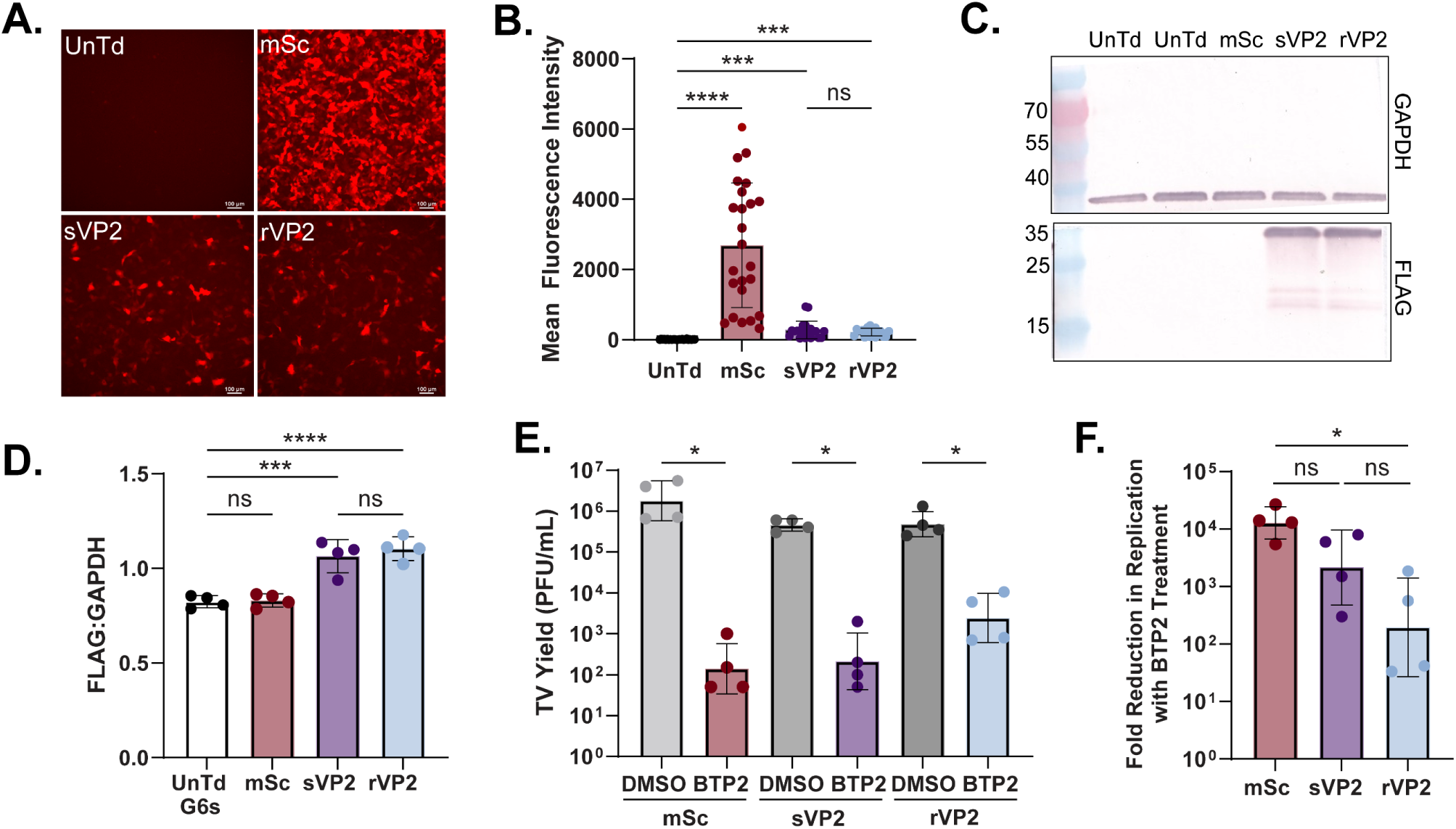
Resistant VP2 partially rescues the replication of BTP2 susceptible TV. **A**) Representative images of mean fluorescence intensity of cells transduced with AAVs expressing VP2 or mScarlet alone. Images were taken at 48 hours post transduction (hpt). Scale bar = 100μM. **B**) Quantitation of mean fluorescence intensity across 4 biological replicates. **C**) Western blot demonstrating expression of FLAG-tagged susceptible or resistant VP2 following AAV transduction. Cell lysates were harvested at 48 hpt. **D**) Quantitation of western blot intensity plotted relative to GAPDH. Each point represents an independent biological replicate. **E**) Plaque assay titration of MA104 cells transduced with AAVs expressing mScarlet or susceptible or resistant VP2, infected with BTP2 susceptible TV (susceptible clone 1) at an MOI 1, and treated with DMSO control or BTP2. Infections were performed 48hpt, and compound treatment was done 1 hpi. **F**) Fold change in virus yield between DMSO and BTP2 treatment conditions from panel E. For all experiments, data was collected across 4 biological replicates. Normality was assessed by Shapiro-Wilk test. TV yield assay data is plotted as geometric mean ± SD, *p<0.05 by unpaired T test. Mean fluorescence intensity data is compared using Kruskal-Wallis with Dunn’s multiple comparisons test. For the fold change and western blot quantitation, *p<0.05 by 1-way ANOVA with Sidak’s multiple comparisons test.

### BTP2 inhibits human norovirus replication

Given that BTP2 inhibited TV replication, we next assessed BTP2 activity against HuNoV using our antiviral pipeline for HIOs[36]. We first evaluated GII.4 Sydney (P16) HuNoV replication in jejunum-derived human intestinal organoids (jHIOs) treated with a dose range of BTP2. We found that 30 µM and 100 µM BTP2 led to a significant reduction in HuNoV replication at 24 hpi, as did the positive control, 50 µM 2’-C-methylcytidine (2-CMC) [36, 37] (Figure 7A). While GII.4 HuNoV is the predominant circulating genotype, we wanted to evaluate the efficacy of BTP2 against other HuNoV strains. We found that 30 µM and 100 µM BTP2 treatment also significantly inhibited GII.3 (P21) replication in jHIOs (Figure 7B). To evaluate the selective index of BTP2 in organoids, we plotted the percent inhibition of viral replication and the percent cytotoxicity in organoids across the BTP2 dose range. We used non-linear regression to determine the half-maximal effective concentration (EC_50_) and half-maximal cytotoxic concentration (CC_50_). By taking the ratio of these metrics, BTP2 had a selective index of 4.74 for GII.4 Sydney (P16) and a selective index of 6.36 for GII.3 (P21) HuNoV (Figure 7C-E). These data suggest that BTP2 has weak antiviral activity against both GII.4 and GII.3 strains of HuNoV.

**Figure 7:**
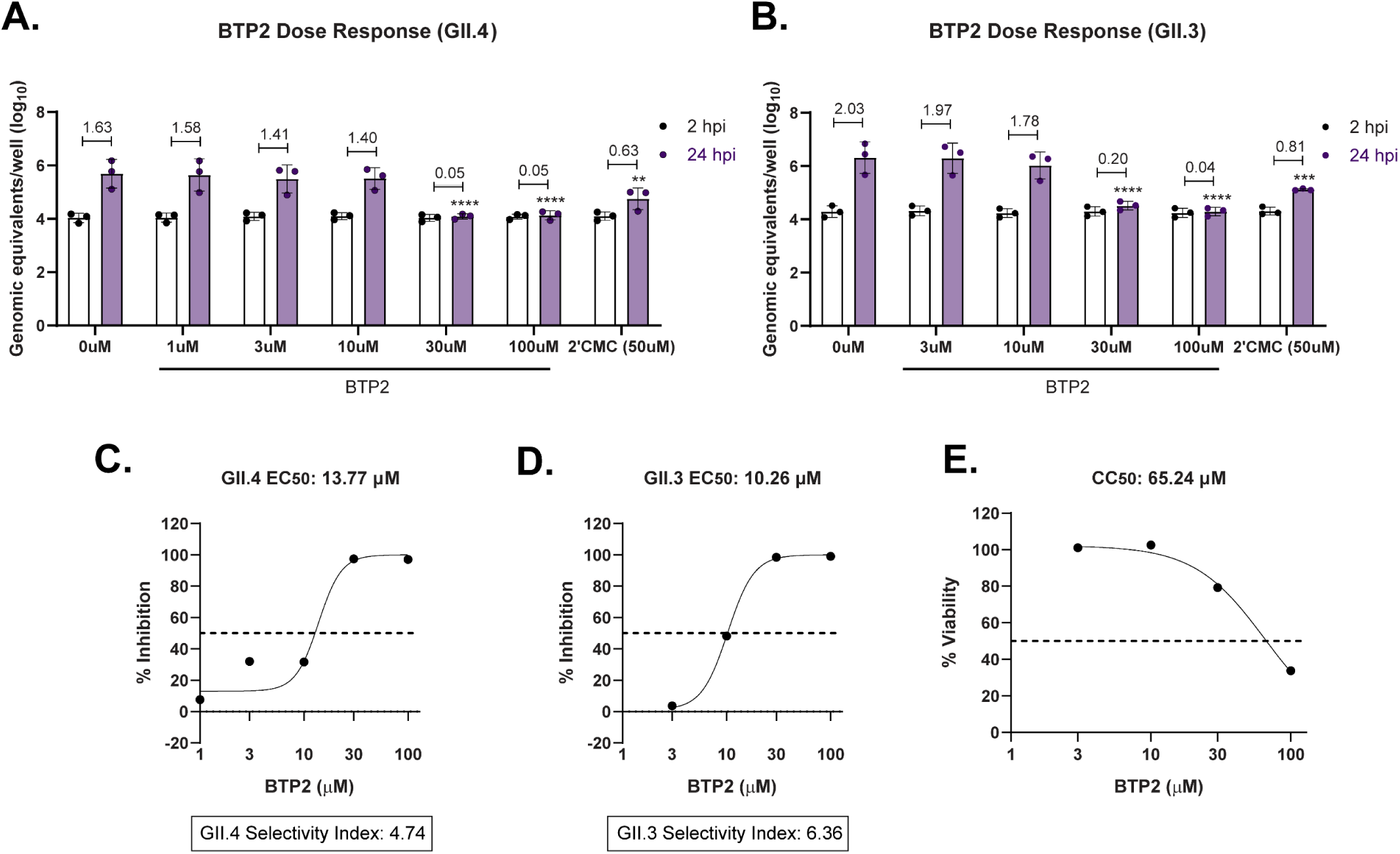
BTP2 inhibits HuNoV replication. **A)** Quantitation of HuNoV genome equivalents in jejunal human intestinal organoids (jHIOs) inoculated with 100 TCID50s of GII.4 virus at 2 and 24 hpi after treatment with a dose range of BTP2 or 50μM 2-CMC. Both compounds were incubated with enteroid monolayers for 1 hr before infection and maintained in the maintenance media until harvest (24 hr). **B**) Quantitation of HuNoV genome equivalents in jHIOs inoculated with 100 TCID50s of GII.3 (P21) virus at 2 and 24 hpi. jHIOs were treated with BTP2 or 2-CMC as described above at the indicated concentrations. **C-D**) Percent inhibition of GII.4 (C) or GII.3 (D) replication across the BTP2 dose range for 3 independent experiments. EC_50_ and CC_50_ values were calculated using non-linear regression and the selective indices were calculated by dividing the CC_50_ by the EC_50_. **E**) Percent viability as measured by LDH assay over a dose range of BTP2. Data are pooled from 5 independent experiments. For HuNoV replication data, ****p<0.0001 by 2-way ANOVA with Dunnett’s multiple comparisons test.

## Discussion

There is a growing body of research that demonstrates a potential use for Orai channel blockers in infectious diseases, as they have shown *in vitro* efficacy in inhibiting virus replication [24]. Viruses across a range of families encode calcium-conducting viral ion channels, which can mediate the release of calcium from the endoplasmic reticulum into the cytosol [38]. In multiple contexts, this has been shown to engage the SOCE pathway. We speculated that Orai would be an attractive broad-spectrum host-directed antiviral therapeutic target. Interestingly, BTP2 was unique in its ability to block TV replication, and this was likely due to direct effects on TV structural proteins rather than on SOCE. Furthermore, BTP2 treatment significantly decreased HuNoV replication. This work identifies the calicivirus structural proteins as viable targets for antiviral development. While future work is needed to confirm that BTP2 acts directly on HuNoV structural proteins, these findings lay the foundation for the characterization of BTP2 as a novel antiviral pharmacophore.

Identifying the TV structural proteins as likely targets of BTP2 allows us to speculate on the potential mechanism by which it can inhibit viral replication. TV VP1 plays an important role in HBGA attachment factor binding [21], and capsid assembly. HuNoV VP1 similarly mediates HBGA binding [39–42] and capsid assembly [43, 44] but is also involved in viral entry [45]. Less is known about the role of VP2 for both viruses, although it has been shown to stabilize HuNoV virus-like particles [46], facilitate genome encapsidation [47], and potentially regulate the activity of the RNA-dependent RNA polymerase [48]. In feline calicivirus and murine norovirus, VP2 has also been shown to be essential for infectivity[49{Ishiyama, 2024 #82]}. These ascribed functions of VP1 and VP2 seem consistent with a dual role for BTP2 in both the early and late stages of infection. Thus, we postulate that BTP2 may be targeting capsid stability, pioneer rounds of translation or genome replication (early) and/or capsid assembly (late) during infection. This is supported by our time-of-addition data, which indicates that treatment with BTP2 either before or after infection significantly reduced TV replication. However, we cannot rule out the possibility that the effect of pre-treatment stems from BTP2 retention within the cells even after media change, allowing BTP2 to exert its antiviral activity later during infection. More directed studies are needed to decipher the mechanisms responsible for the reduction in replication observed with BTP2 treatment.

Our serial plaque isolation experiments yielded three BTP2-resistant clones with the VP1(I380M) mutation, and one clone (BTP2-resistant clone 3) with no amino acid changes in VP2. Interestingly, resistant clone 3 exhibited moderate BTP2 resistance compared to the other two BTP2-resistant clones, both of which had additional mutations in VP2. Thus, we aimed to confirm the roles of both VP1 and VP2 in mediating BTP2 sensitivity. Our reverse genetics studies indicate that the VP1 resistance-associated amino acid change is indeed resistance-conferring. The rescue of TV replication in the presence of BTP2 with the VP1 I380M mutation alone also suggests that other viral mutations or features (e.g., 5’ or 3’ UTRs) are not necessary for BTP2 resistance. It is unclear if VP2 plays an accessory role in drug susceptibility, as virus with only VP2 resistance-associated amino acid changes was still susceptible to BTP2 treatment. It is possible that higher concentrations of BTP2 are needed to detect differences in susceptibility between susceptible and VP2 only rescued viruses, or VP1 only and dual VP1 and VP2 viruses, however these findings will be complicated by increased cytotoxicity of BTP2 above the 100µM concentration. Ectopic overexpression of the resistant VP2 was also not sufficient to fully rescue the replication of clonally susceptible TV in the presence of BTP2. However, a partial rescue was seen compared to expression of mScarlet alone. This suggests that VP2 is not sufficient for, but may contribute to or enhance, BTP2 resistance. Further work using TV reverse genetics to systematically test the various VP2 mutations with and without the VP1 I380M mutation will be critical to dissect the importance of these changes. While we find it most likely that BTP2 acts directly upon VP1 and/or VP2, there is also the possibility that BTP2 alters a host pathway in a manner that inhibits susceptible TV but the amino acid changes in VP1 and/or VP2 in the resistant TV circumvents viral dependence on these pathways. Thus, further investigation of how BTP2 inhibits TV replication and how the amino acid changes in VP1 and VP2 confer resistance to BTP2 will provide new mechanistic insights into calicivirus replication strategies in addition to identifying new functional targets for antiviral drug development.

It was surprising to find that TV was unique in its susceptibility to BTP2 among the other recovirus strains tested. While none of the TV resistance-associated amino acid changes are completely conserved in the FT7, FT285, or Mo/TG30 sequences, the overall amino acid identity between the structural proteins of these viruses is quite low (∼75% for VP1 and between 60-68% for VP2). Similarly, given that HuNoV and TV VP1 and VP2 share only ∼30% and ∼27% amino acid identity respectively, it is challenging to assess whether the mechanism of action of BTP2 is conserved across these viruses using sequencing data alone. Thus, further work is needed to verify that the HuNoV structural proteins mediate BTP2 susceptibility in the same way that we hypothesize to be true for TV. Identifying the defining features of BTP2 susceptibility in HuNoV would be particularly important, as it may allow us to predict which circulating strains of virus will be most potently inhibited. Additionally, these findings would have important implications for understanding the likelihood of, and potential barriers to, a virus acquiring *de novo* BTP2 resistance. Finally, identifying the precise mechanism of BTP2 resistance would facilitate optimized drug design to impede resistance development.

Our results are consistent with previous work, which identifies BTP2 as a known potent inhibitor of SOCE in both T cells [50, 51] and MA104 cells [26]. While the mechanism of BTP2 inhibition of SOCE has yet to be fully characterized, T cell studies indicate that BTP2 acts extracellularly to inhibit Orai channel activity [50]. Our time-of-addition studies are most consistent with the ability of BTP2 to permeate MA104 cells after prolonged treatment. Given that SOCE assays are done with acute treatment, it is likely that BTP2 maintains both an extracellular function in inhibiting Orai and an intracellular function in inhibiting viral replication. This work is also not the first to examine the potential effects of BTP2 on virus replication. We have previously demonstrated that BTP2 reduces rotavirus replication [26]. While it is tempting to speculate that this may be due to SOCE-independent effects, STIM1 knockdown also reduces rotavirus replication [24], which we did not observe to be the case for TV. Thus, we cannot fully rule out an on-target, SOCE-mediated role for BTP2 in rotavirus infection. Other cellular off-target effects of BTP2 have also been previously identified, including inhibition of the actin reorganizing protein, drebrin [52], inhibition of TRPC channels [53], and potentiation of the nonselective cation channel, TRPM4 [54]. The contributions of these off-target effects to the ascribed inhibitory action of BTP2 against either SOCE or viral replication must be fully explored to understand the BTP2 antiviral mechanism.

Given both the established on- and off-target effects of BTP2, understanding its value as a candidate therapeutic requires attention to how BTP2 may be tolerated in patients. While new SOCE channel inhibitors are in development for a range of human diseases, few have been FDA-approved due to lack of specificity and potential toxicity. However, one known Orai channel blocker, CM4620 (Auxora), has reached phase II clinical trials for the treatment of acute pancreatitis [55] and COVID-19 pneumonia [56]. Additionally, a number of FDA-approved compounds, though not originally developed as Orai channel blockers, have since been shown to significantly reduce SOCE *in vitro*[57]. These include leflunomide, teriflunomide, lansoprazole, tolvaptan, and roflumilast. Both leflunomide and teriflunomide demonstrated inhibitory activity against SOCE at their clinically relevant doses [57]. Of note, these compounds were screened in part based on their structural similarity to BTP2. BTP2 has also been evaluated in preclinical mouse, rat, and guinea pig models of autoinflammatory disease after acute oral or intraperitoneal treatment and is well tolerated up to 30 mg/kg when administered per-orally [51, 58–62]. In addition to standard animal models, organoids are becoming an increasingly important model for preclinical compound validation. BTP2 treatment of *ex vivo* human intestinal epithelial organoids at 1 µM concentration did not significantly alter cell viability, cell differentiation, or barrier function [63]. In addition, our work demonstrates that >80% human jejunal organoid viability is maintained with 30 µM BTP2 treatment.

Excitingly, our studies demonstrate that BTP2 has antiviral activity against GII.4 and GII.3 strains of HuNoV. To better contextualize its efficacy, we compared the antiviral activity of BTP2 to a positive control, 2-CMC, and determined its selective index. We used 2-CMC as the positive control because it is a nucleoside analog originally developed as a hepatitis C virus (HCV) antiviral and significantly inhibits MNV replication [64] and HuNoV in the Norwalk replicon system [65], B cells [66], and HIOs [36, 37, 67]. Currently, 2-CMC is one of the most selective and potent anti-norovirus compounds identified to date; however, its prodrug, valopicitabine, failed phase II clinical trials in the treatment of HCV due to gastrointestinal toxicity [68]. Thus, while 2-CMC is not a viable antiviral candidate, it represents a strict comparator for evaluating compound efficacy *in vitro*. Selective indices are an important pharmacologic metric used to account for both the potency of an antiviral therapeutic (EC_50_) and its related toxicity (CC_50_)[36]. We found that the selective indices of BTP2 for GII.4 Sydney (P16) and GII.3 (P21) HuNoV are 4.74 and 6.36, respectively. While there is no strict, universal cutoff for therapeutic screens, selective indices greater than or equal to 10 have been used to indicate potential clinical utility. In context, 2-CMC has shown a selective index greater than 31 against GII.4 Sydney in HIOs [36]. Thus, while BTP2 itself may not have optimal selective activity, the strong inhibition of TV replication and broad, albeit lesser antiviral activity against HuNoV strains support the idea that BTP2 may be a promising starting pharmacophore that can be used to optimize or screen other candidate norovirus antiviral therapeutics.

Together, these studies define an important SOCE-independent role of the Orai channel blocker, BTP2, in inhibiting virus replication. In addition, our work demonstrates that BTP2 is a new, optimizable therapeutic for HuNoV infection. More broadly, we establish the utility of targeting norovirus structural proteins in antiviral development and present a platform whereby candidate antiviral compounds can be screened in surrogate virus systems, like TV and/or other recoviruses, and validated with HuNoV infection in organoids, facilitating the discovery of other novel antiviral therapies.

## Materials and Methods

### Cell lines and viruses

MA104 African Green Monkey kidney epithelial cells were engineered to stably express the cytosolic calcium indicator, GCaMP6s, by lentivirus transduction as previously described [69]. MA104G6s STIM1 knockout cells were cloned by limiting dilution and validated in an SOCE assay and via western blot (Supplemental Figure 1). Cells were incubated at 37°C in 5% CO_2_ and maintained in Dulbecco’s Modified Eagle Medium (DMEM) supplemented with 10% Fetal Bovine Serum (FBS) and Antibiotic-Antimycotic (Gibco, 1X final concentration). TV was generated by serial passage in MA104 cells. All experiments were performed with virus that has been passaged at least 19 times. TV stocks were generated by infecting MA104 cells at MOI 0.01 and harvesting 48 hpi (when ∼95% CPE was observed). Virus was titrated by plaque assay on MA104 cells (see below). The FT7 and FT285 virus was provided to our lab by Dr. Tibor Farkas [45] and Mo/TG30 was provided by Dr. David Wang [19]. GII.4(P31) and GII.3(P21) HuNoV stocks were prepared as 10% stool filtrates as described previously [7].

### Human intestinal organoids

Jejunal human intestinal organoids (HIOs) were obtained from the Gastrointestinal Experimental Model Systems Core at the Texas Medical Center Digestive Diseases Center. HIOs suspended in Matrigel were cultured in “WRNE” culture medium containing Wnt3a, R-spondin-3, Noggin, EGF and passaged every 7 days. HIOs were dispersed into a single cell suspension and plated as monolayers prior to differentiation for HuNoV infection studies. HIO differentiation was achieved by placing monolayers in Intesticult differentiation medium (STEMCELL Technologies) for 5 days before infection [6, 70].

### Compounds

BTP2 was purchased from Millipore Sigma (CAS 223499-30-7), Ro2959 from MedChemExpress (CAS 2309172-44-7), GSK-7975A from AOBIOUS (CAS 1253186-56-9), and Synta66 from Millipore Sigma (835904-51-3). Ro2959 was used at a final concentration of 5 µM, and BTP2, GSK, and Synta66 were used at a final concentration of 10 µM. In experiments where BTP2 was added to the plaque assay overlay, a final concentration of 5 µM was used to avoid cytotoxicity with >48 h of incubation. All compounds were dissolved in DMSO, which served as our vehicle control for all experiments. Thapsigargin was purchased from Thermo Scientific (CAS 67526-95-8) and used at a final concentration of 1 µM.

### SOCE Assays

MA104G6s cells were seeded on µClear 96-well imaging plates (Greiner) at a density of 50,000 cells per well in 10% FBS DMEM maintenance medium. Once confluent, the cells were incubated with Orai channel inhibitors or DMSO for 20 minutes before imaging. After pre-treatment, cell monolayers were washed with calcium-free Hanks’ Balanced Salt Solution (HBSS) and equilibrated in the epifluorescence microscope live cell imaging chamber. Imaging was performed with a Nikon TiE inverted microscope with a SPECTRAX LED light source (Lumencor) using a 10X Plan Apo (NA 0.30) objective. Cells were imaged in a 2 sec interval for 1-2 minutes to establish a baseline fluorescence before perfusion of 1 µM thapsigargin solution in calcium-free HBSS, again containing Orai channel inhibitors or DMSO. Following an additional 6-8 minutes of continuous imaging, the perfusion solution was then changed to calcium containing HBSS (2 mM calcium) with inhibitor or control. The cells were imaged for an additional 4-5 minutes, after which the imaging run was completed. SOCE traces were generated by plotting the GCaMP6s fluorescence per field-of-view (FOV) overtime. The maximum GCaMP6s delta fluorescence was calculated by subtracting the baseline normalized fluorescence of the FOV from the maximum fluorescence intensity following 2mM calcium addition and plotting this relative to the DMSO control.

### Cytotoxicity Assays

Cytotoxicity was measured in MA104 cells or J2 jejunal human intestinal organoids (jHIOs) using the Promega CytoTox 96 Non-Radioactive Cytotoxicity assay, according to the manufacturer instructions with minor modifications for HIOs[71]. All experimental and control compounds were incubated with the monolayers at the indicated concentrations for 24 hours. jHIOs were differentiated for 5 days prior to addition of the compounds, and all jHIO supernatants and controls were diluted 1:10 prior to running the assay as previously described [71].

### Virus Infections

MA104 monolayers were incubated in serum-free DMEM for 24 hrs prior to infection. TV inoculum was prepared at the indicated MOI in serum-free DMEM medium and absorbed on cell monolayers for 1 hour at 37°C in 5% CO_2_. Following incubation, virus inoculum was removed, and cell monolayers were washed with PBS before adding the maintenance medium. HuNoV infections were performed on differentiated HIO monolayers in a 96-well plate pre-treated for 1 hr with vehicle control, BTP2, or 50µM 2-CMC. Cells were infected with 100 TCID_50_ of GII.3 or GII.4 HuNoV diluted in CMGF-(Advanced DMEM F12, 1X Glutamax, 10mM HEPES, 100 U/mL penicillin/streptomycin). After a 1-hour inoculum incubation, cells were washed with CMGF- and placed in Intesticult (Stem Cell Technologies) differentiation medium containing BTP2 at the indicated concentrations or 50µM 2-CMC. At 24 hpi, cell lysates and supernatants were harvested for lactate dehydrogenase (LDH) cytotoxicity assay or RT-qPCR.

### Virus Yield Assays

MA104cells were seeded in 24-well tissue culture-treated plates at a density of 100,000 cells per well and grown to confluency. Cells were infected with TV at an MOI of 1 as described above. Following inoculum incubation, cells were placed in medium containing Orai channel inhibitors or DMSO vehicle control. At 24 hours post-infection, cells were harvested by three freeze-thaw cycles and virus was quantitated by plaque assay. For plaque assays, MA104 cells were seeded in a 6-well tissue culture plate at a density of 500,000 cells per well and grown to confluency. Cells were inoculated with 10-fold serial dilutions of infected lysates as described above. Following infection, cell monolayers were overlaid with a 1.2% microcrystalline cellulose solution, made by combining equal parts 2.4% microcrystalline cellulose (Avicel) with 2X DMEM, and supplementing with 0.1 mg/mL DEAE dextran. Plaque assays were harvested 72 hpi and monolayers were stained with crystal violet to visualize plaques.

### Virus Growth Curves

MA104 cells were seeded in 24-well tissue culture-treated plates at a density of 100,000 cells per well and grown to confluency. Cells were infected with TV (parental or serial plaque isolated) at an MOI of 1 as described above. Following inoculum incubation, cells were placed in medium containing Orai channel inhibitors or DMSO vehicle control at the indicated time points. 1, 6, 12, and 24 hours after infection, cells were harvested by three freeze-thaw cycles and virus was quantitated by plaque assay.

### Western Blot

MA104 cells were mock inoculated or infected with BTP2 susceptible TV (susceptible clone 2) at an MOI of 1 as described above and treated with either DMSO or 10µM BTP2 at 1, 6, or 10 hpi. Cell lysates were harvested at 24hpi in RIPA buffer (50mM Tris base, 150mM NaCl, 1% NP-40, 0.5% sodium deoxycholate, and 0.1% sodium dodecyl sulfate). Samples were homogenized using a biopolymer column (QiaShredder, Qiagen), combined with SDS-PAGE sample buffer, and heated at 100°C for 10 minutes prior to loading. Proteins were separated on 4-20% Tris-glycine SDS-PAGE gels (BioRad) and transferred onto a nitrocellulose membrane. Blocking was performed with 10% non-fat dry milk in PBS. Primary antibodies (Rabbit αTV VP1, synthesized by ABClonal; mouse αGAPDH, Southern Biosciences; FLAG) were diluted in 0.5% blocking solution, used at a concentration of 1:2000, and incubated overnight at room temperature. Secondary antibodies (Goat αRabbit-AP, Goat αMouse-AP) were also diluted in 0.5% blocking solution, used at a concentration of 1:2000 and were incubated for 2 hours at room temperature. Membranes were washed with 0.5% blocking solution and developed with an alkaline phosphatase detection solution containing 50mM Tris, 3mM MgCl_2_, 0.1 mg/mL p-nitro blue tetrazolium chloride, and 0.05 mg/mL 5-bromo-4-chloro-3-indolyl phosphate. Blots were imaged using a document scanner (Canon) and band intensity was quantified by converting blots to 8-bit grayscale images in ImageJ, performing uniform background subtraction, inverting the pixel intensity, and taking the ratio of the sum intensity of each band to that of the respective GAPDH loading control. Quantitation is from 3 blots run with lysate from three independent infections.

### TV BTP2 Plaque Assays

MA104 cells were seeded in 6-well plates at a density of 500,000 cells/well and grown to confluency. Cells were inoculated with 10-fold serial dilutions of virus stock (3×10^7^ PFU/mL-3×10^8^ PFU/mL, depending on MA104 passage number) as described above. 3 mL of 1.2% microcrystalline cellulose overlay containing 5µM BTP2 or an equivalent volume of DMSO was added to each of the wells. Plaque assays were harvested 72 hpi and monolayers were stained with 1X crystal violet to visualize plaques. Plaque size was quantitated by measuring the diameter of well-isolated plaques.

### Solid Overlay Plaque Assays and Plaque Picking

TV solid overlay plaque assays were performed as described above, with the modification that a 1.2% agarose and neutral red overlay were used in place of the microcrystalline cellulose overlay. The first SeaKem agarose overlay was made by combining equal parts 1.2% SeaKem agarose (SeaKem LE agarose, Lonza) solution and 2X DMEM medium, supplemented with DEAE dextran and either 5µM BTP2 or DMSO. 48 hours after the first overlay, a second overlay containing equal parts 1.2% SeaKem agarose solution and 2X DMEM medium, supplemented with 0.05% neutral red was added to each of the wells. 24 hours after the second overlay was added, plaques were picked by harvesting the overlay plug and adding it to 1 mL of serum free DMEM. Plaque picks were then used to inoculate MA104 cells to generate stocks in the presence of DMSO or 10µM BTP2 for downstream yield, plaque, and CPE inhibition assays.

### Cytopathic Effect Virus Inhibition Assay

The CPE inhibition assay was performed as previously described [72]. Briefly, MA104 cells were grown to confluency in 24-well tissue culture plates. Once confluent, cell monolayers were inoculated with TV at an MOI 10 as described above. Infected monolayers were treated with DMSO or BTP2 at the indicated concentrations and incubated for 24 hours, after which maintenance medium was removed and the cells were washed with PBS. Monolayers were then stained with 1X crystal violet for 10 minutes with rocking. The unbound stain was removed with PBS wash and monolayers were allowed to dry at room temperature. Bound crystal violet stain was eluted with 33% acetic acid (400 µL per well), which was incubated for 10 minutes at room temperature, with rocking. From each well, 100uL of crystal violet elution were transferred to a 96-well plate and absorbance was measured at 630nm (BioTek ELx808 spectrophotometer). The inhibition rated (%) was calculated with the following equation: inhibition rate = [(OD_inhibitor-treated infected cells_ - OD _DMSO-treated infected cells_)/(OD _DMSO-treated uninfected cells_ - OD _DMSO-treated infected cells_)] x 100.

### Virus Drug Passaging

Passage 1 TV was generated by infecting MA104 cell monolayers with TV at an MOI 1 as described above. After inoculation, monolayers were incubated in serum-free medium containing DMSO vehicle control or 10µM BTP2. Infected lysate was harvested by three freeze-thaw cycles at 24 hpi. Passage 2-4 TV was generated by infecting MA104 monolayers in a 24-well plate with 200uL of lysate from the previous passage. The lysate was allowed to absorb for 3 hours at 37°C in 5% CO_2._ After the inoculation period, the lysate was removed, monolayers were washed with PBS, and medium was replaced with serum-free DMEM containing DMSO control or 10µM BTP2. Infected lysate was harvested at ∼32 hpi by three freeze-thaw cycles.

### TV VP2 Transduction and Co-infection

AAV vectors expressing mCherry and 3X FLAG-tagged TV VP2 or mScarlet alone were commercially synthesized and packaged (VectorBuilder). For transduction, MA106cells were seeded in a 24-well plate at a density of 150,000 cells per well. One day post-seeding, cell monolayers were transduced with susceptible or resistant VP2 (clone 1) at an MOI of 25,000 genome copies per cell or mScarlet at an MOI of 1,000 genome copies per well. 48 hours post transduction, cells were imaged to assess transduction efficiency and subsequently harvested in RIPA buffer or infected with TV (susceptible clone 1) at an MOI 1. Infected cells were treated with either DMSO or 10µM BTP2 1 hpi. After 24 hours, infected cell lysates were harvested and titrated via plaque assay, as described above.

### TV Sequencing

RNA was isolated from MA104 cells infected with the serial plaque isolated virus at MOI 3 in the presence of either DMSO or BTP2 using a viral RNA kit (Zymogen), according to the manufacturer’s instructions. RNA yield and quality was assessed on a spectrophotometer (Nanodrop, Thermo Scientific). cDNA synthesis and genome amplification were done in 2 segments, an ORF1 and an ORF2,3 segment, following manufacture protocol (One-Step RT-PCR kit, Takara). PCR product size was verified by gel electrophoresis. Primers used are listed in Table 1. PCR products were column purified (Promega Wizard PCR Cleanup Kit) and sequenced by Sanger sequencing (Azenta). Sequencing primers are listed in Table 2. Sequencing contigs were assembled and aligned using SnapGene (ver. 7.2.0) software.

**Table 1.**
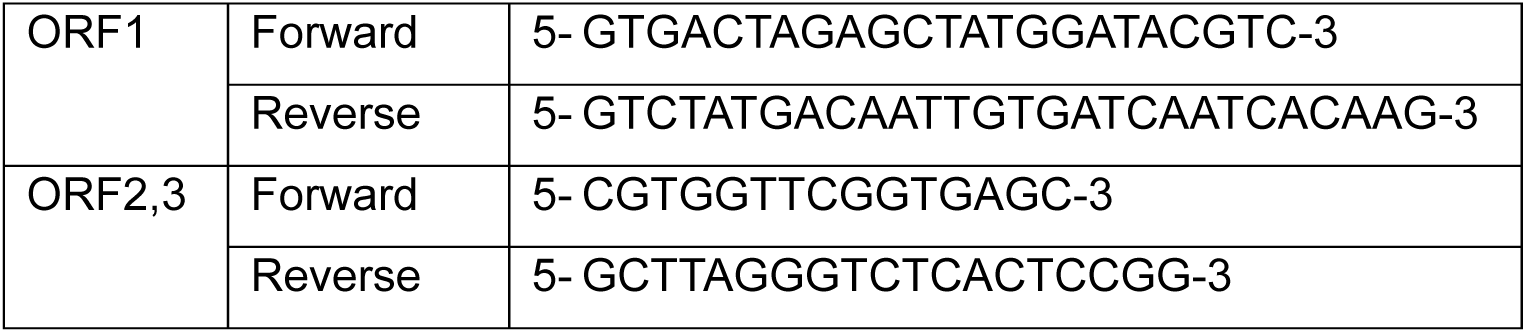
Primers for TV one-step RT-PCR.

**Table 2.**
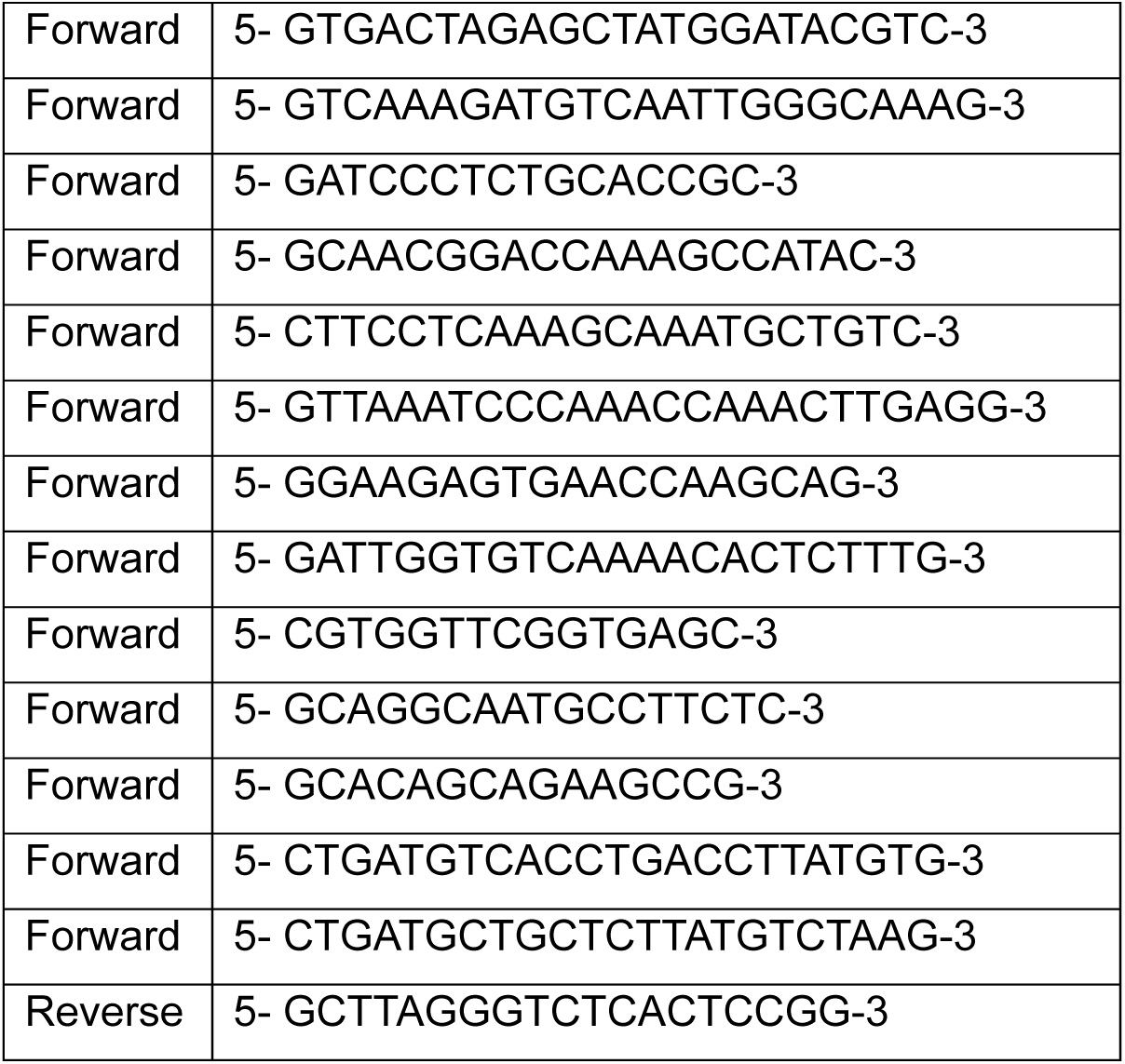
Primers used for Sanger Sequencing.

### Quantitating TV Genome Equivalents through RT-qPCR

MA104 cells were seeded in a 24-well plate and grown to confluency. Cells were infected with TV (susceptible clone 2) at MOI 1 and treated with BTP2 1 hour before inoculation, during inoculation, or at 1, 6, or 10 hours post inoculation. At 24 hpi, cells were harvested, and RNA was isolated using column-based extraction (Zymogen viral RNA kit) per the manufacturer’s instructions. RNA yield and quality was assessed on a spectrophotometer (Nanodrop, Thermo Scientific). cDNA was synthesized from 800ng of RNA (SensiFast cDNA synthesis kit, Bioline USA Inc). qPCR was performed by dye incorporation (SYBR green, Invitrogen) and fluorescence quantification (QuantStudio, Applied Biosystems). Each sample was run in technical duplicate. Viral RNA copy number was calculated based on the CT value of the sample compared to a standard curve of total *in vitro* transcribed RNA with known copy numbers. Primer pairs for detection of TV at the 5’ end of the genome were forward: 5-GTCAAAGATGTCAATTGGGCAAAG-3 and reverse: 5-CCCAAGGCACCCAAAACC-3.

### Quantitating HuNoV Genome Equivalents through RT-qPCR

RNA was extracted from infected HIO monolayers using the MagMAX-96 viral RNA isolation kit on the Kingfisher Flex machine (Thermo Scientific). cDNA synthesis and PCR amplification was performed using the qScript XLT One-Step RT-qPCR ToughMix kit (Quantabio) with the following primer pair and probe sets: COG2R/QNIF2d/QNIFS. Viral RNA copy number was calculated based on the CT value of the sample compared to a standard curve based on recombinant GII.4 Houston virus (HOV) RNA transcripts[7].

### FRET protease assays

Tulane protease and GII.4 Sydney protease were expressed in E. coli with N-terminal 6xHis-TELSAM fusion tag with a HRV3C protease cleavage site and purified by Ni-NTA chromatography, tag cleavage and removal, and size exclusion chromatography. 2.5X solutions of protease (Tulane: 5µM, GII.4 Sydney: 2.5µM) and 2X solutions of FRET substrate (Glu(EDANS)-GDYELQGPEDLA-Lys(Dabcyl), 40µM) were prepared by dilution of more concentrated stocks with assay buffer (10mM HEPES, 30% glycerol, 10mM DTT, 0.1% CHAPS, pH 8.0). 10X stocks of BTP2 of various concentrations were diluted from 10mM BTP2 stock with DMSO. Assays were run with 100ul reaction volumes in 96-well black NBS plates (Corning 3991) at 37°C and read with the FlexStation 3 (Molecular Devices) with 90-second measurement intervals (excitation 340nm, emission 490nm, filter 475nm).

### Cell-free Polymerase Assay

GII.4 RdRp activity was measured using a real-time fluorescence-based assay, which uses SYTO9, a fluorescence dye that specifically binds dsRNA but not ssRNA template molecules. Reactions were performed in individual wells of black 96-well flat-bottom plates (costar). The standard reaction contained GII.4 RdRp (1 μM), 20 mM Tris-HCl pH 7.5, 5 mM MgCl2, 2.5 mM MnCl2, 40 μg/mL polyC, 5 U RNAseOUT (invitrogen) and 0.25 μM SYTO9 (Sigma-Aldrich). The reaction was initiated by the addition of 300 μM GTP and the fluorescence was recorded every 5 min over 180 min at 37 °C using a plate reader FlexStation3 (Molecular devices). To investigate the effect of BPT2 on the polymerization activity of RdRp, 1 μM RdRp was incubated with 100 μM BPT2 at room temperature for 1 hour and the RdRp activity was measured using the above-mentioned protocol.

### Tulane virus Reverse Genetics

cDNA plasmids encoding the TV genome with the BTP2 susceptible or resistant-associated VP1 and/or VP2 sequences downstream of a CBh RNA PolII and K1E phage promoter were synthesized commercially (Vector Builder). MA104 cells were transfected with 7.5ug of the TV genomic DNA plasmid and 2.5ug of C3P3, a T7 polymerase-African swine fever virus capping enzyme fusion protein [73]. Transfections were performed with the TransIT-LT1 transfection reagent at a concentration of 2µL per 1µg of DNA, per the manufacturer instructions. CPE was observed ∼72 hours post transfection (hpt), upon which cells were harvested by three freeze/thaw cycles. The rescued virus was passaged twice in MA104 cells, titrated by plaque assay and utilized for BTP2 yield and dose range studies as described above. Viral sequences were confirmed by sanger sequencing (Azenta).

## Acknowledgements

This work was supported by National Institutes of Health Grants R01DK115507 and R01AI158683 (PI: J.M. Hyser), and P30DK56338 (PI: M.K. Estes) that supports the Texas Medical Center Digestive Diseases Center and the Gastrointestinal Experimental Model Systems (GEMS) core. Trainee support for F.J.S. was provided by NIH grants F31DK132942 (PI: F.J. Scribano). Trainee support for J.T.G was provided by NIH grants F30DK131828 and Histochemical Society Graduate Medical Trainee and Graduate Student Cornerstone Grant (PI: J.T. Gebert). Trainee support for K.A.E. was provided by NIH grants F32DK130288 and Histochemical Society Postdoctoral Keystone Grant (PI: K.A. Engevik).

## Supplemental Figures

**Supplemental Figure 1:**
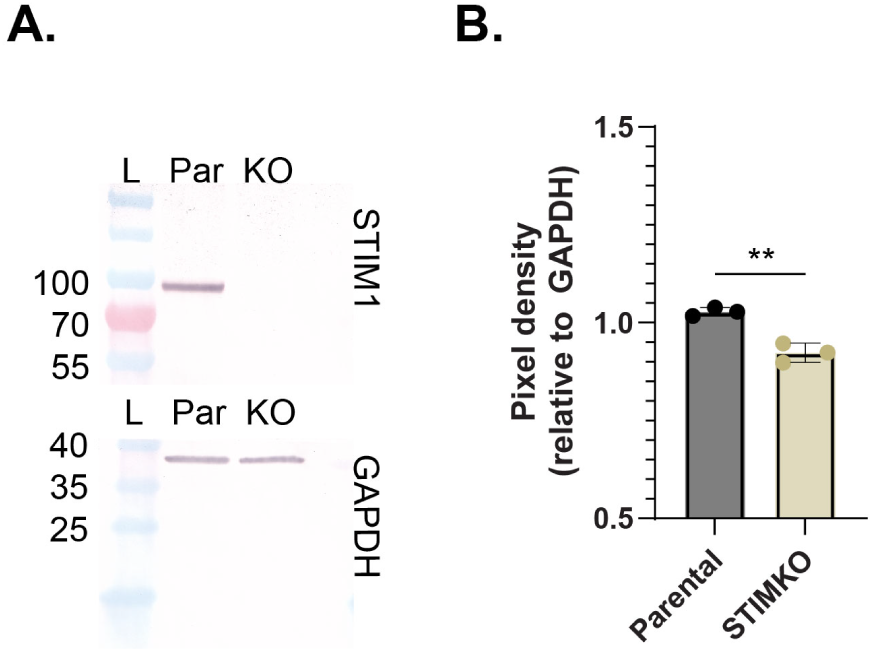
MA104G6s STIM1KO cells have undetectable STIM1 protein expression. **A**) STIM1 protein expression (top) detected by Western in MA104G6s parental (Par) or STIM1 knockout (KO) cells. GAPDH (bottom) served as the loading control. **B**) Quantitation of band intensity plotted relative to GAPDH. Each point represents an independent biological replicate. **p<0.01 by unpaired T test.

**Supplemental Figure 2:**
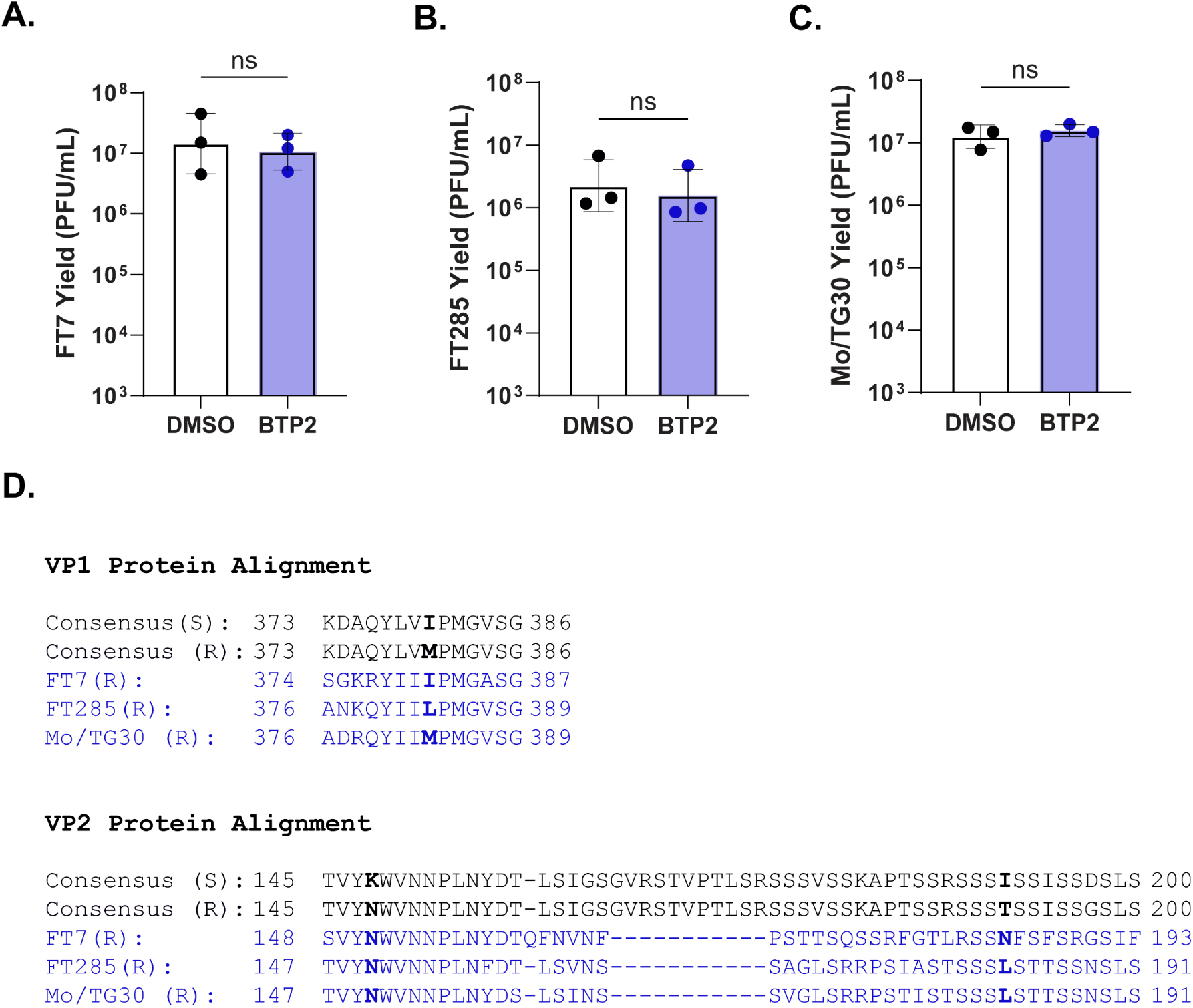
BTP2 does not inhibit the replication of other recoviruses. **A-C**) Recovirus FT7 (A), FT285 (B), and Mo/TG30 (C) yield in PFU/mL at 24hpi with DMSO (white) or 10uM BTP2 (blue) treatment at 1hpi. Experiments were performed with three biological repeats of two averaged technical replicates and data are plotted as geometric mean ± SD. For all experiments, normality was assessed by Shapiro-Wilk test and analyzed by unpaired T test. **D**) Alignment of the consensus BTP2 susceptible (S) and resistant (R) TV VP1 and VP2 sequences and the Recovirus FT7 (GenBank: KC662368.1), FT285 (GenBank: KC662366.1), and Mo/TG30 (GenBank: OQ184950.1) VP1 and VP2 sequences (blue).

**Supplemental Figure 3:**
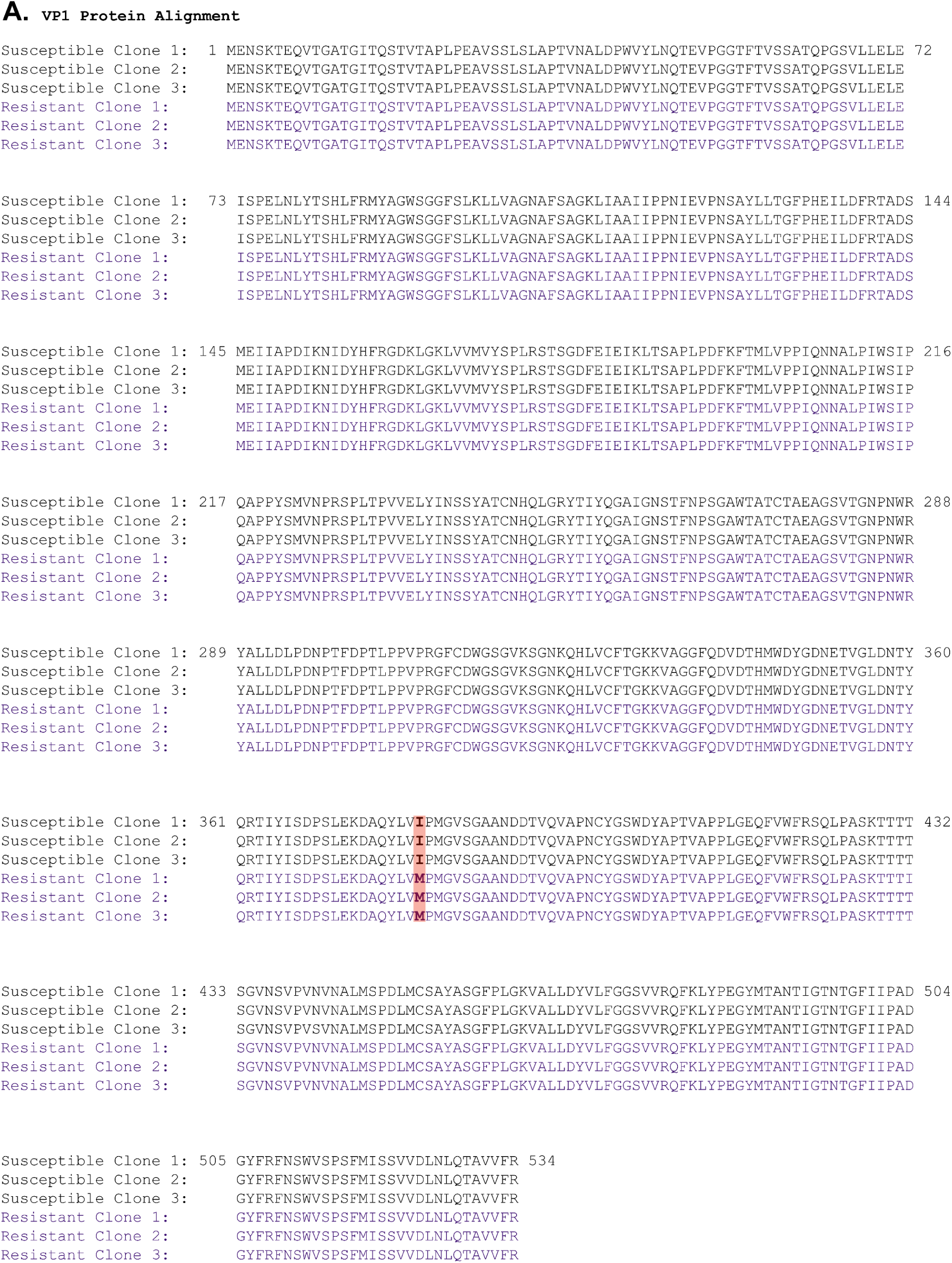
VP1 amino acid alignment. **A**) VP1 amino acid alignment between 3 BTP2 susceptible and 3 BTP2 resistant variants. Conserved amino acid differences are highlighted in red.

**Supplemental Figure 4:**
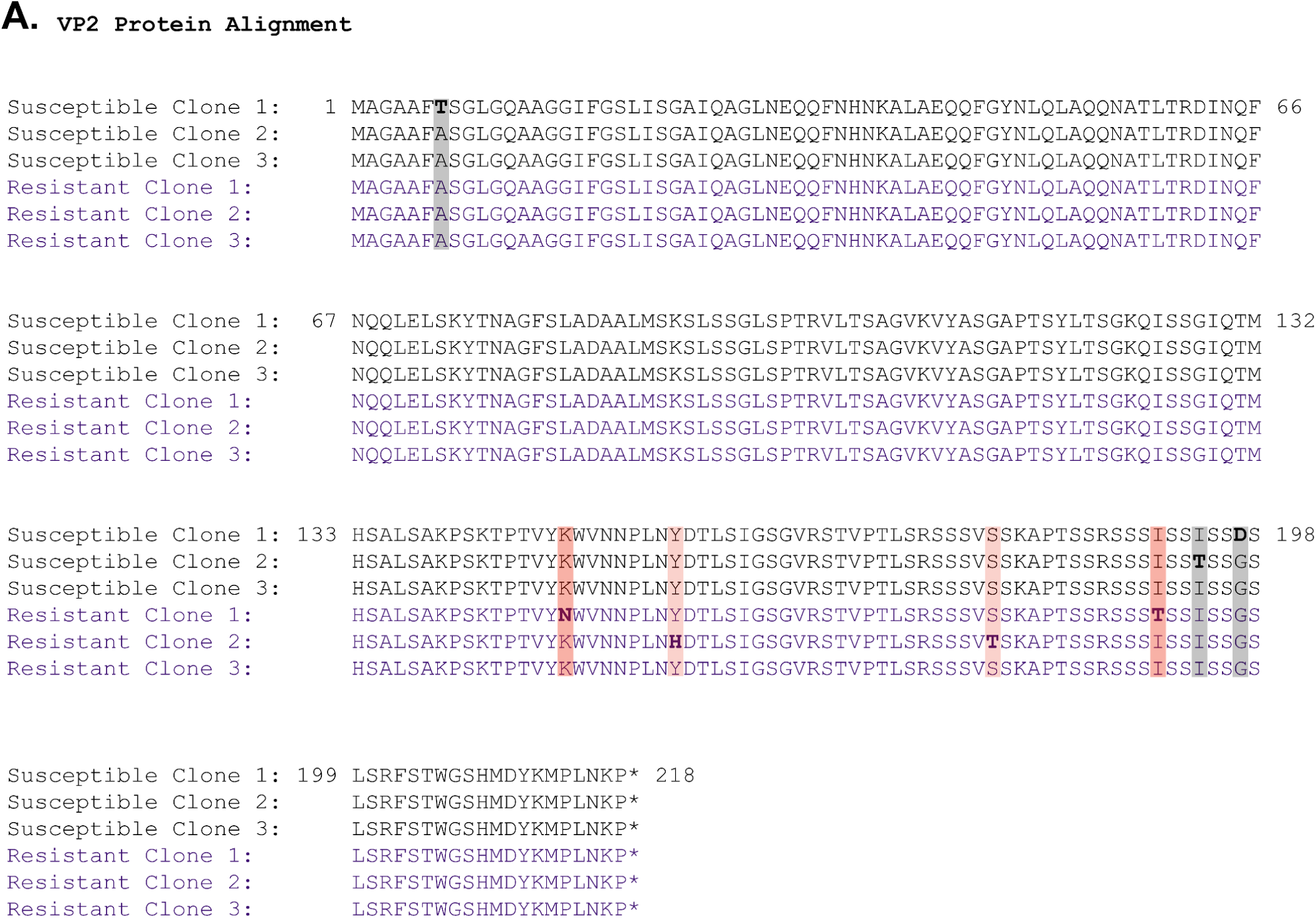
VP2 amino acid alignment. **A**) VP2 amino acid alignment between 3 BTP2 susceptible and 3 BTP2 resistant variants. Amino acids that differ from the consensus sequence in the resistant (red) or susceptible (grey) clones are highlighted.

**Supplemental Figure 5:**
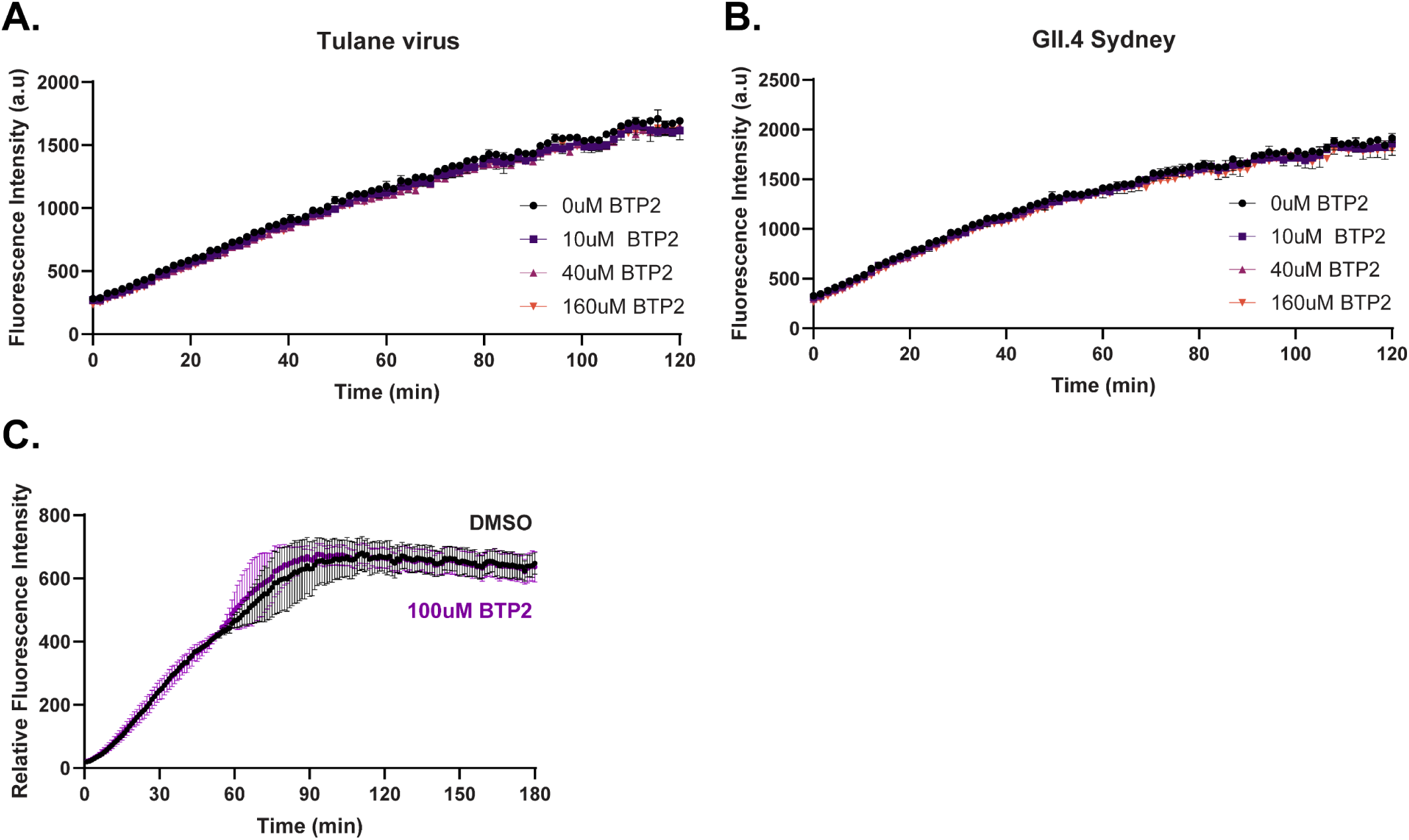
BTP2 does not Inhibit TV or HuNoV Protease or Polymerase activity. **A-B**) Cell free FRET assay measuring TV (A) or GII.4 Sydney HuNoV (B) protease activity in the absence (black) or presence of BTP2 at the 10μM (purple), 40μM (pink), or 160μM (orange) concentration. C) GII.4 RDRP activity in the presence of DMSO (black) or 100μM BTP2 (purple).

**Supplemental Figure 6:**
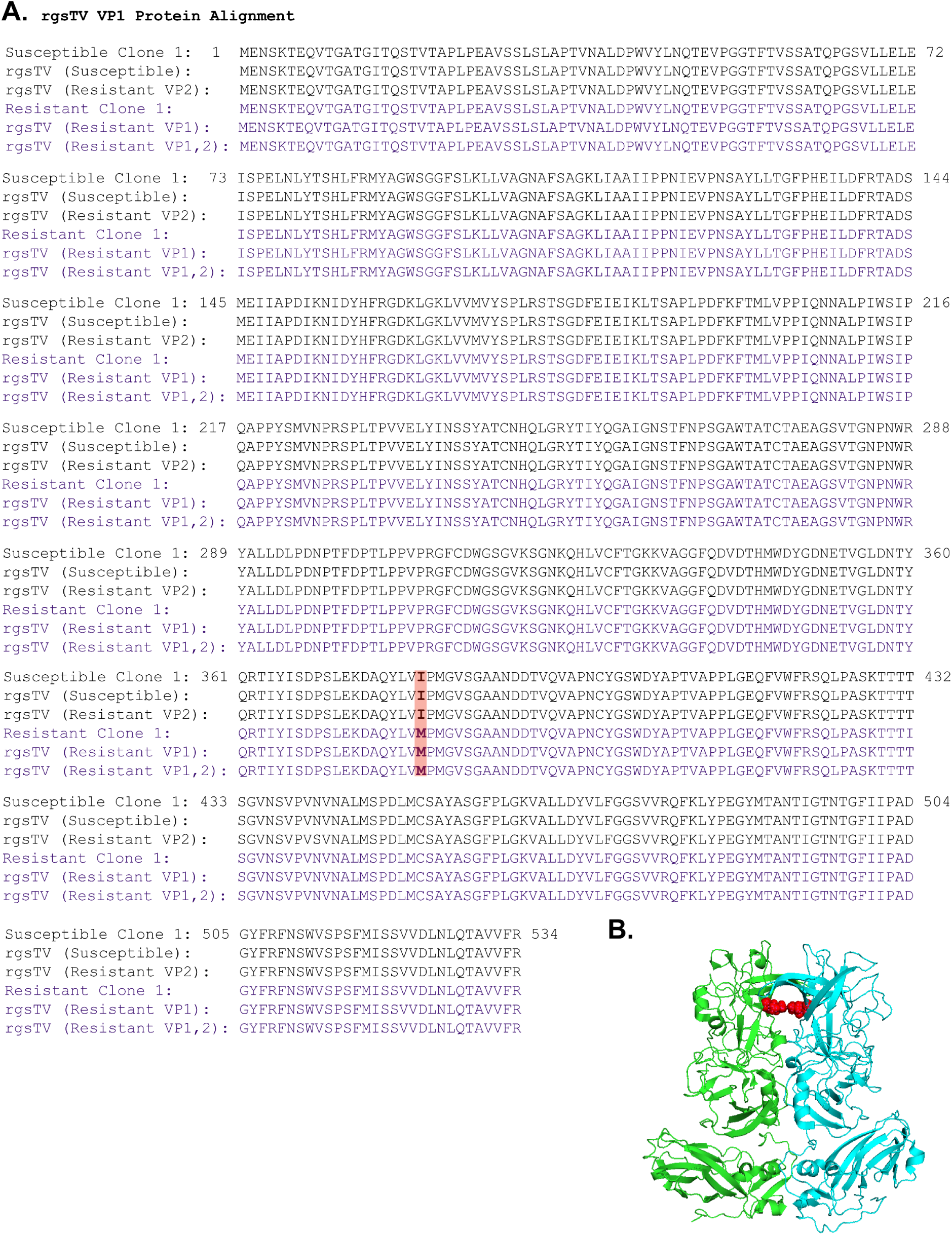
Reverse genetics TV VP1 amino acid alignment. **A**) VP1 amino acid alignment of the clone 1 BTP2 susceptible and clone 1 BTP2 resistant TV sequences with the reverse genetics Tulane viruses. **B**) Mapping of the isoleucine at position 380 (red) on the VP1 dimer.

**Supplemental Figure 7:**
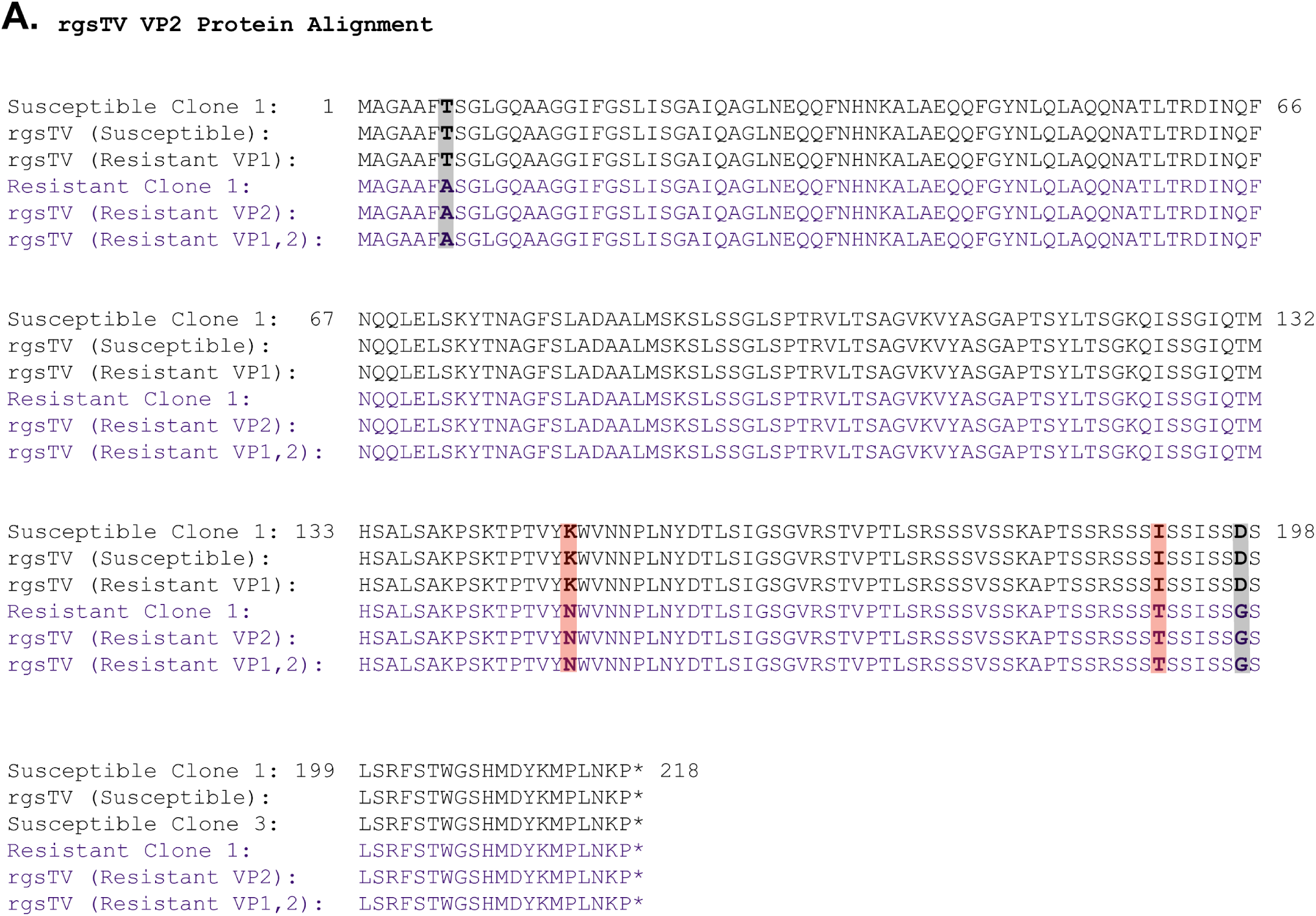
Reverse genetics TV VP2 amino acid alignment. **A**) VP2 amino acid alignment of the clone 1 BTP2 susceptible and clone 1 BTP2 resistant TV sequences with the reverse genetics Tulane viruses.

